# ALK signalling primes the DNA damage response sensitizing ALK-driven neuroblastoma to therapeutic ATR inhibition

**DOI:** 10.1101/2023.08.30.555570

**Authors:** Marcus Borenäs, Ganesh Umapathy, Dan E. Lind, Wei-Yun Lai, Jikui Guan, Joel Johansson, Eva Jennische, Alexander Schmidt, Yeshwant Kurhe, Jonatan L. Gabre, Agata Aniszewska, Anneli Strömberg, Mats Bemark, Michael N. Hall, Jimmy Van den Eynden, Bengt Hallberg, Ruth H. Palmer

## Abstract

High-risk neuroblastoma (NB) is a significant clinical challenge. MYCN and ALK, which are often involved in high-risk NB, lead to increased replication stress in cancer cells, suggesting therapeutic strategies. We previously identified an ATR/ALK inhibitor (ATRi/ALKi) combination as such a strategy in two independent genetically modified mouse NB models. Here, we identify an underlying molecular mechanism, in which ALK signalling leads to phosphorylation of ATR and CHK1, supporting an effective DNA damage response. The importance of ALK inhibition is supported by mouse data, in which ATRi monotreatment resulted in a robust initial response, but subsequent relapse, in contrast to a 14-day ALKi/ATRi combination treatment that resulted in a robust and sustained response. Finally, we show that the remarkable response to the 14-day combined ATR/ALK inhibition protocol reflects a robust differentiation response, reprogramming tumour cells to a neuronal/Schwann cell lineage identity. Our results identify a unique ability of ATR inhibition to trigger neuroblastoma differentiation and underscore the importance of further exploring combined ALK/ATR inhibition in NB, particularly in high-risk patient groups with oncogene-induced replication stress.

## Introduction

High-risk neuroblastoma (NB) is a childhood cancer that currently presents a clinical challenge, reflected in the fact that NB accounts for 15% of all paediatric tumour related deaths^1^. Although aggressive NB initially responds to treatment, later relapses exhibit poor prognosis and survival rates of around 35%^2,3^. For this reason, extensive efforts are being made to identify more effective therapeutic options. The relative paucity of mutations in NB has made it challenging to identify targetable therapeutic options. Genes with increased somatic mutation frequencies in NB include *ALK*, *PTPN11*, *ATRX* and *NRAS*, where ALK can be therapeutically targeted by small molecule tyrosine kinase inhibitors (TKIs)^4–9^. While not mutated, amplification of the *MYCN* transcription factor is observed in approximately 20% of NB and is an important predictive factor that is currently therapeutically intractable^10,11^. Although lacking in mutations, NB exhibits a high number of somatic chromosomal lesions, including deletion of regions of chromosome 1p, 11q, and gain of 2p, 17q, as well as aneuploidy, all of which provide important prognostic information^6^.

Increased genome instability, which is commonly seen in tumours, leads to the engagement of DNA damage sensor systems^12^. The DNA damage response (DDR) employs members of the PI3-kinase-related protein kinase (PIKK) family, such as the ataxia telangiectasia and Rad3-related (ATR) and ataxia-telangiectasia mutated (ATM), to ensure DNA integrity^13^. ATR also has an important role in cell survival in response to replication stress, by preventing replication origin firing and reducing the number of active forks, maintaining a stability of stalled replication forks, facilitating repair and promoting replication restart^13,14^. ATR was recently identified as a potent therapeutic target in preclinical models of NB^15–18^ and we demonstrated that combined inhibition of ATR (with elimusertib) and ALK (with lorlatinib) results in a complete ablation of tumours in ALK-driven-NB GEMMs. While we identified ATR as a downstream target of ALK signalling in NB cells^15^, it is unclear what consequence ALK signalling activity has for ATR and the DDR. To date the underlying molecular mechanisms involved in the dramatic responses to ALK/ATR inhibition in ALK-driven models of NB^15^ have been elusive.

Here we demonstrate the superiority of ALKi/ATRi combination therapy over monotherapy and show that this combined effect is explained by ALK priming the DDR through phosphorylation of ATR on Ser 435 and CHK1 on Ser 280. We further identify a robust neuronal and Schwann cell differentiation response in tumours treated with ATR inhibitors. Taken together, these results strongly motivate the continued exploration of ATR/ALK inhibition as a potentially therapeutically effective approach in NB.

## Results

### Combined ATR/ALK inhibition is more effective than monotreatment in ALK-driven mouse NB models

In previous work we reported a robust and sustained complete response to combinatorial ALK and ATR inhibitor treatment in two ALK-driven GEMM models of NB^15^. To better understand the mechanism behind the combination response, we compared a 14-day ATRi monotreatment with the ALKi/ATRi combinatorial regime in *Alk-F1178S;Th-MYCN*^19^ mouse NB tumours. Mice harboring tumours were treated with a 14-day regime that either combined elimusertib together with the ALK TKI lorlatinib (3 days elimusertib 25 mg/kg twice daily, 4 days lorlatinib 10 mg/kg twice daily, 3 days combination, 4 days lorlatinib) or employed elimusertib alone (3 days elimusertib 25 mg/kg twice daily, 4 days vehicle twice daily, 3 days elimusertib 25 mg/kg, 4 days vehicle) (Fig. 1A-B). Tumour volumes were monitored with ultrasound at 4, 7, 11 and 14 days during the 14-day treatment. As previously, all mice tolerated the treatment regime with no noticeable side effects^15^. Further, no tumours were detected at day 14 after combined ALKi/ATRi treatment (Fig. 1C). Elimusertib monotherapy also resulted in a significant reduction in tumour size relative to control but did not result in a complete resolution of tumour material at 14 days. Mice were observed over time without any further therapeutic interventions and were monitored regularly with ultrasound for tumour development. Remarkably, 14-day combinatorial treatment (elimusertib together with lorlatinib), resulted in a sustained complete response as we have previously reported^15^. In contrast, monotreatment with elimusertib resulted in tumour relapse, with 50% of treated mice relapsing within 43 days of treatment compared with 0% relapse at this time in mice treated with the elimusertib/lorlatinib combination (log-rank (Mantel-Cox) test, *P* < 0.05; log-rank test; Fig. 1D).

**Figure 1.**
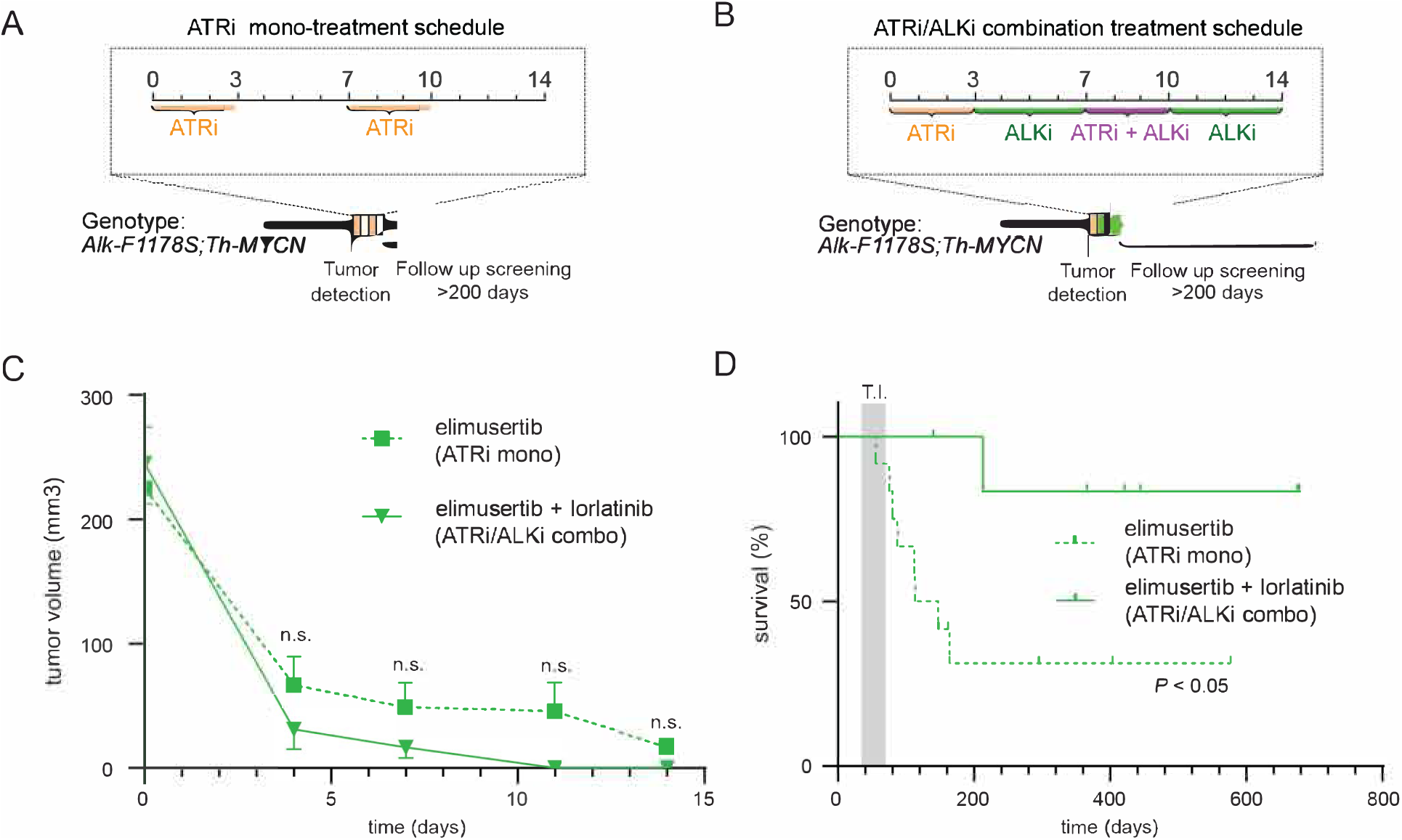
ALK inhibition enhances tumour response and progression free survival in mice treated with ATR inhibitors. **A.** Schematic overview of the monotreatment regimen for Alk/MYCN-driven GEMM tumours. In brief, mice were treated with 25 mg/kg ATRi (elimusertib) for 3 days, after this a 4 day pause from treatment was then followed by an additional 3 days of elimusertib. **B.** In the combination regimen the initial monotreatment with ATRi was followed by 10 mg/kg lorlatinib for 11 days, with the addition of ATRi at days 8–10. All treatments were given orally twice daily. **C.** Tumour volume measured by ultrasound over 14 days. Data is presented as mean +/- SEM. n.s. = not significant. **D.** Survival (from birth) of *Alk-F1178S;Th-MYCN* treated with elimusertib monotreatment (n=12) compared with elimusertib/ lorlatinib combination treatment (n=7). *P* < 0.05; log-rank (Mantel-Cox) test. Shaded area indicates tumour incidence (T.I.) range.

### ALK signalling primes the DNA damage response in NB cells

Our earlier analysis of ALK TKI treatment in NB cells identified changes in the phosphoproteome downstream of ALK that included ATR^20^. To reveal the effect of ALK signalling on ATR and the DDR, we performed a detailed phosphoproteomics analysis of NB cells (NB1) stimulated with ALKAL2 ligand. This confirmed ATR as a phosphorylation target downstream of ALK, with ALKAL2 stimulation leading to an increase in phosphorylation of Ser435 (log2FC = 1.754; *P =* 1.4×10^-5^), that has been reported to be a growth factor stimulated ATR site that regulates ATR activity^21^. In addition to ATR phosphorylation, ALK stimulation also induced phosphorylation of the ATR target CHK1 at Ser280 (log2FC = 1.56; *P* = 2.6 x10^-4^). Of note, this ALKAL2 stimulated CHK1 phosphorylation was not at the ATR S/TQ target Ser345 that is paradoxically increased in response to CHK1 inhibition by LY2603618^22^ (Fig. 2A). In line with these findings, inhibition with the ALK TKI lorlatinib resulted in decreased CHK1 Ser280 phosphorylation (log2FC = -1.31; *P =* 6.8×10^-6^). We confirmed phosphorylation of CHK1 on Ser280 in response to ALKAL2 stimulation in two independent NB cell lines, CLB-BAR and NB1 (Fig. 2A). Phosphorylation of CHK1 S280 was independent of ATR, as addition of elimusertib did not block phosphorylation in response to ALK activation. Conversely, inhibition of ALK with lorlatinib in ALK-driven CLB-BAR, CLB-GE and CLB-GAR NB cell lines resulted in decreased phosphorylation of pCHK1 S280 (Fig. 2B). It has been reported that phosphorylation of CHK1 on Ser280 regulates CHK1 localisation^21^. To investigate this, NB cells were stimulated with ALKAL2 and cytoplasmic and nuclear fractions were immunoblotted for pCHK1 S280. A robust increase in pCHK1 S280 on ALK stimulation was detected in both cytoplasmic and nuclear fractions, although no significant difference in the total amount of CHK1 in the nuclear fraction was detected (Fig. 2C). Since the phosphorylation of CHK1 on S280 has been reported to accelerate the CHK1 activation process and prime the DDR, we investigated the effect of ALK inhibition on the DDR. ALK-driven CLB-BAR, CLB-GE and CLB-GAR NB cells were treated with lorlatinib for 24 h and phosphorylation of H2A.X on S139 monitored as a readout of DNA damage. Indeed, ALK inhibition resulted in an increase of pH2A.X that could be observed at 24 h in all three cell lines (Fig. 2D). These results suggest that ALK inhibition decreases the cellular DDR, eventually resulting in DNA damage. This was confirmed by analysis of 276 genes that have been associated with the DDR^23^ (these genes are further referred to as the DDR signature) in previously published RNA-Seq datasets from different NB cell lines treated with lorlatinib^19,20^. Lorlatinib treatment for 6 h or more resulted in a significant reduction of the DDR signature in ALK-driven CLB-BAR, CLB-GE and NB1 but not in the ALK wild type IMR32 and SKNAS NB cell lines (Fig. 2E). Similar DDR reductions were observed for other ALK inhibitors (e.g., crizotinib, entrectinib) and other ALK-driven NB cell lines (e.g., CLB-GAR) for which RNA-Seq data are publicly available (Fig. S1). Taken together, our data suggest that ALK signalling modulates the DDR in ALK-driven NB cells. To test this hypothesis, we next treated ALK-driven CLB-BAR, CLB-GE and CLB-GAR NB cells with lorlatinib (30nM) or etoposide (500nM), either alone or in combination for 24 h (Fig. 2F). Treatment with etoposide alone resulted in an increase in pH2A.X, that was further increased on ALK inhibition, confirming that ALK signalling enhances the DDR in these cells, and suggesting a molecular mechanism by which ALK inhibition is able to increase the efficacy of ATR treatment.

**Figure 2.**
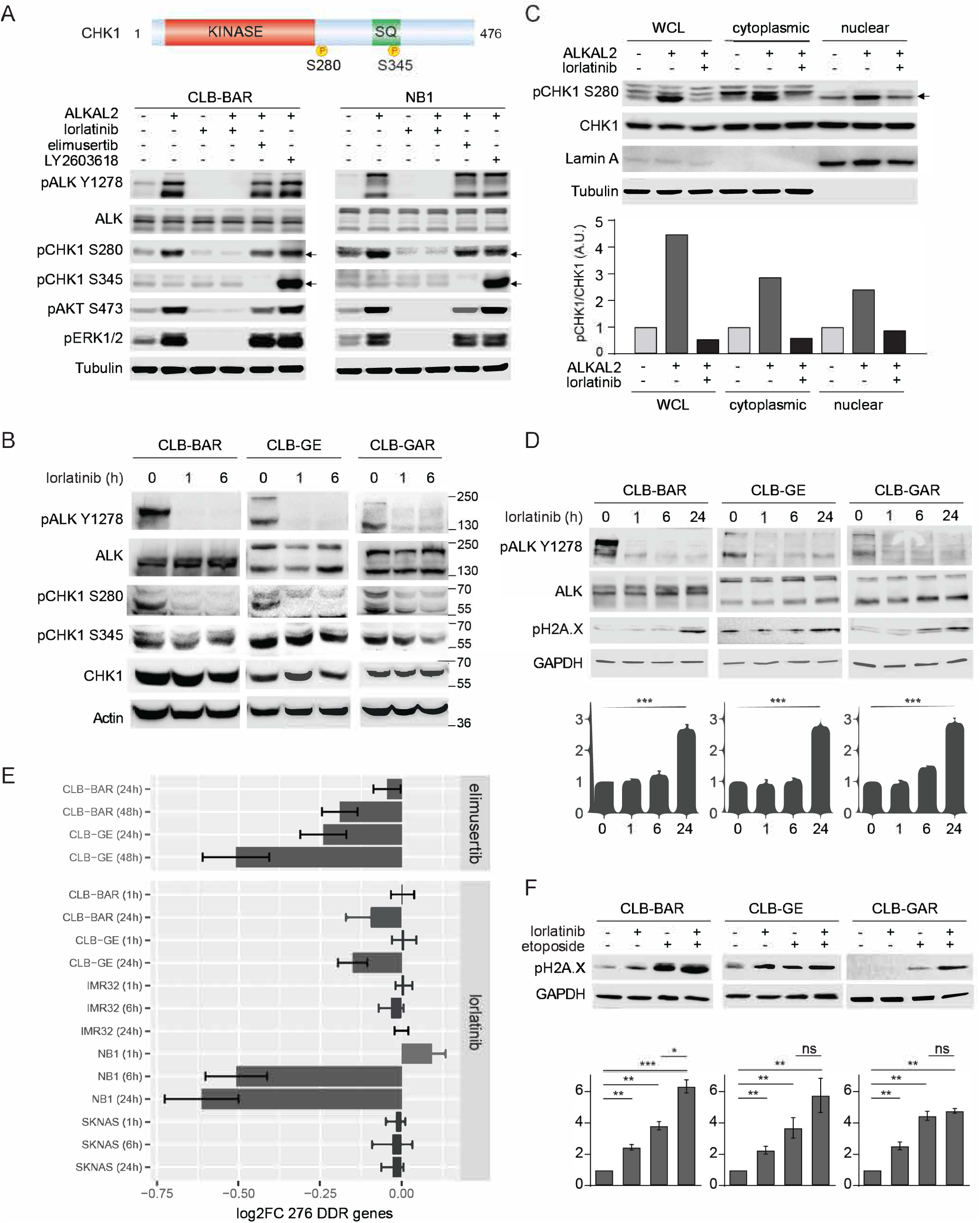
ALK signalling activity drives phosphorylation of CHK1 on S280. **A.** ALK-driven CLB-BAR and NB1 cells were treated with different inhibitors (2 h) or ALKAL2 ligand (0.5 h) alone or in combination as indicated. Inhibitors employed were the ALK inhibitor lorlatinib (20 nM), the ATR inhibitor elimusertib (100 nM), and the CHK1 inhibitor LY2603618 (1 μM). Cell lysates were immunoblotted for pALK (Y1278), ALK, pCHK1 (S280), pCHK1 (S345), pAKT (S473), pERK1/2 (T202/Y204) and Tubulin. The serine sites S280 (downstream of ALK) and S345 (downstream of ATR) are highlighted schematically in CHK1 (top panel). Kinase domain (in red) and regulatory SQ domain (SQ, in green). Arrowheads indicate pCHK1 (S280) and pCHK1 (S345). **B.** ALK-positive CLB-BAR, CLB-GE, and CLB-GAR NB cells were treated with lorlatinib (30 nM) for 0, 1, and 6 h. Cell lysates were immunoblotted for pALK (Y1278), ALK, pCHK1 (S280), pCHK1 (S345), CHK1, and Actin. **C.** Whole cell lysates, cytoplasmic extracts and nuclear extracts of CLB-BAR cells treated with ALKAL2 ligand alone or in combination with lorlatinib were immunoblotted for pCHK1 (S280), CHK1, the nuclear protein marker Lamin A and cytoplasmic protein marker β-tubulin. Arrowhead indicates pCHK1 (S280). Quantification of pCHK1/CHK1 ratios normalized to controls is shown below. **D.** ALK-positive CLB-BAR, CLB-GE, and CLB-GAR NB cells were treated with lorlatinib (30 nM) at different time points, as indicated. Cell lysates were immunoblotted for ALK, pALK (Y1278), CHK1, pCHK1 (S280), pCHK1 (S345), and p-H2A.X (S139). GAPDH was employed as loading control. n = 3 biologically independent experiments. Unpaired t test; \*\*\**P*<0.001. **E.** Bar plot showing RNA-Seq based log2FC values (mean ± 95% confidence interval) of 276 genes involved in the DNA Damage Response (DDR) for different NB cell lines and drug treatments as indicated. Data derived from 5 previously published studies^15,19,20,83,84^ with DDR genes as defined by Knijnenburg *et al.*^23^. **F.** ALK-positive CLB-BAR, CLB-GE, and CLB-GAR NB cells were treated with lorlatinib (30 nM) or etoposide (500 nM), either alone or in combination for 24 h. DNA damage was monitored by immunoblotting of p-H2A.X (S139). GAPDH was employed as loading control. n = 3 biologically independent experiments. Unpaired t test; ns, not significant; \**P*<0.05; \*\**P*<0.01; \*\*\**P*<0.001.

### Elimusertib is more potent than ceralasertib in NB cell lines

Paediatric clinical Phase I/II trials with elimusertib have recently been initiated (clintrials.gov; NCT05071209), but no clinical data have been reported. Another ATR inhibitor, ceralasertib, was recently employed in combination with the Aurora-A kinase inhibitor MLN8237 (alisertib) with responses in MYCN-driven NB GEMM models^16^. We therefore compared ceralasertib with elimusertib in our experimental setup. While ALK-driven CLB-BAR and CLB-GE cell lines were sensitive to both ATR inhibitors, elimusertib displayed higher potency (IC50s of 67.51 ± 10.23 nM for CLB-BAR and 49.68 ± 8.42 nM for CLB-GE, in agreement with our previous findings^15^) than ceralasertib (IC50s of 480.1 ± 9.56 nM for CLB-BAR and 813 ± 11.06 nM for CLB-GE) (Fig. 3A). These IC50 values agree with previously published analysis of ceralasertib in NB cell lines^18^. To confirm these findings, we compared treatment of CLB-BAR and CLB-GE NB cells with either 50 nM elimusertib, 50 nM ceralasertib or 1 μM ceralasertib for 24 h, immunoblotting for pATR, ATR, pATM, ATM, pCHK1, CHK1, PARP/cl.PARP, p53 and pH2A.X (Fig. 3B). As expected, 50 nM elimusertib inhibited phosphorylation of ATR S428 and its downstream target CHK1 (pCHK1 S345) and resulted in increased levels of cleaved PARP, p53 and pH2A.X. In contrast, 50 nM ceralasertib was insufficient for ATR inhibition and 1 μM was required to effectively inhibit ATR signalling and the DDR.

**Figure 3.**
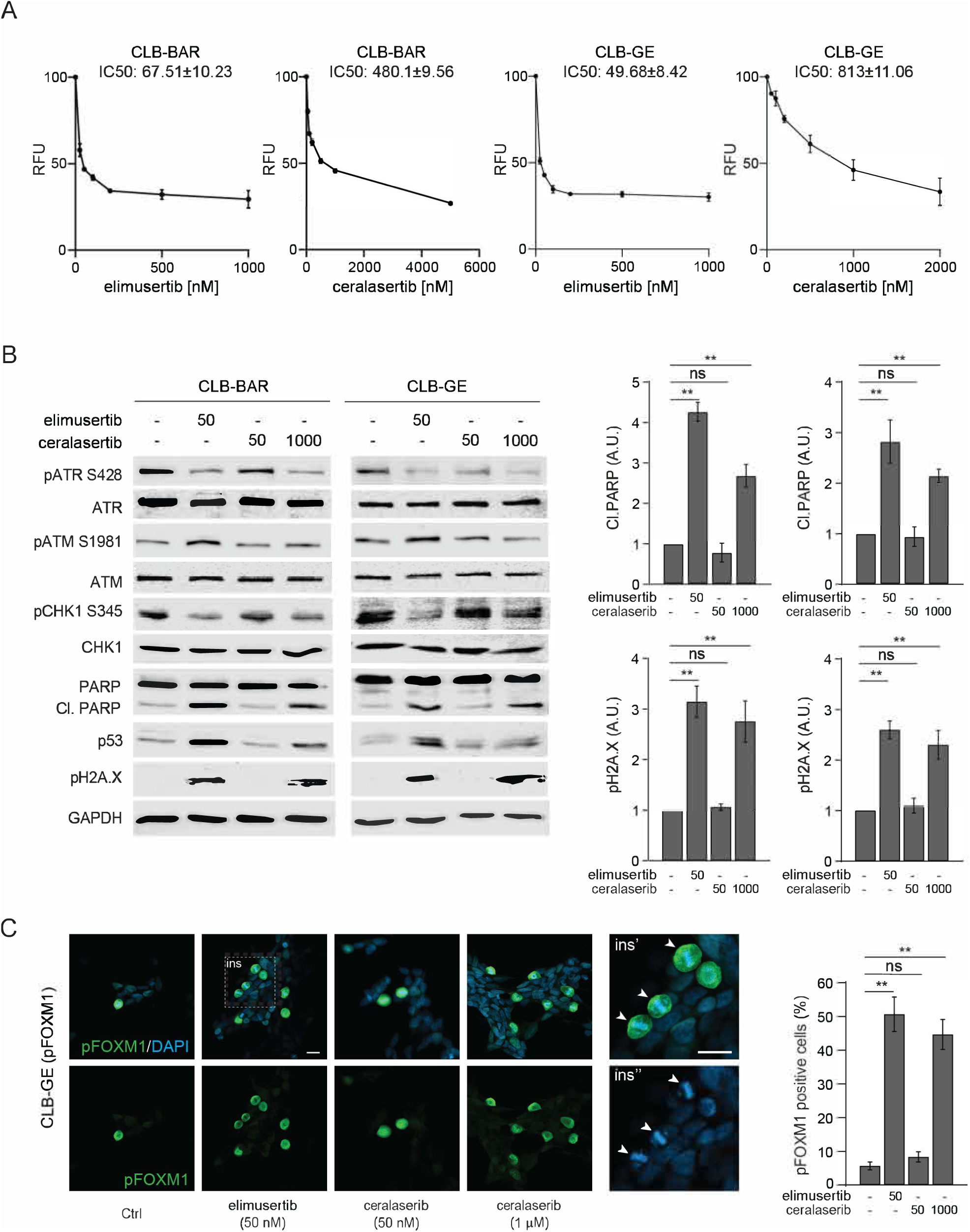
ALK-positive NB cell lines exhibit differential sensitivity to elimusertib and ceralasertib. **A.** CLB-BAR and CLB-GE cell viability in response to increasing concentrations of elimusertib and ceralasertib. Data shown as mean ± SE of fold relative fluorescence units (RFU) relative to untreated cells from three independent experiments. **B.** ALK positive CLB-BAR and CLB-GE NB cells were treated for 24 h with elimusertib or ceralasertib as indicated. Cell lysates were immunoblotted for pATR (S428), ATR, pATM (S1981), ATM, pCHK1 (S345), CHK1, PARP/cl.PARP (quantified at right), p53 and pH2A.X (S139, quantified at right) antibodies. GAPDH was employed as loading control. **C.** CLB-GE cells were synchronized by thymidine block for 24 h followed by treatment with elimusertib or ceralasertib as indicated. Cells were stained for pFOXM1 (green) and DAPI (blue). At right, quantitation of cells positive for pFOXM1 (T600).

High levels of ceralasertib were also required to perturb ATR regulation of the S/G2 checkpoint that controls mitotic entry^24^ (Fig. 3C). The ATR targets pFOXM1 (T600) and pCHK1 (S345) were monitored in CLB-GE cells synchronised by thymidine block and levels of pFOXM1 and pCHK1 monitored in the presence or absence of elimusertib (50 nM) or ceralasertib (50 nM or 1 μM) for 6 h after thymidine release (Fig. 3C, Fig. S2). In agreement with previous findings^15^, enhanced pFOXM1, prominent in metaphase, was observed at earlier time points in CLB-GE cells released from thymidine block in response to ATR inhibition by either 50 nM elimusertib or 1 μM ceralasertib, but not at 50 nM ceralasertib (Fig. 3C). Thus, similar effects were noted for both elimusertib and ceralasertib, although the specific inhibition by ceralsertib demands a 20-fold higher amount compared with elimusertib.

### Phosphoproteomics profiling of ceralasertib in NB cells

We previously characterized the effect of elimusertib mediated ATR inhibition on the NB cell phosphoproteome, identifying a strong compensatory activation of ATM that resulted in an S/TQ target phosphorylation in response to treatment^15^. Although a previous phosphoanalysis by Schlam-Babayov and colleagues employed AZ20, from which ceralasertib is derived^25^, there is no information on the effect of either AZ20 or ceralasertib on the NB phosphoproteome.

To better understand the different effects of elimusertib and ceralasertib, we compared the phosphoproteome of cell treated with the two inhibitors individually. CLB-BAR cells were treated with either 50 nM ceralasertib, 1 μM ceralasertib or 50 nM elimusertib for 6 h after release from thymidine block. A total number of 11,026 phosphosites (9,655 Ser, 1,351 Thr and 20 Tyr) were identified in 3,244 different proteins (Table S1). Differential phosphorylation (DP) was observed in 618 sites (521 phosphorylated, 97 dephosphorylated) from 368 proteins in response to elimusertib treatment (log2FC threshold 0.3 at 5% FDR; Fig. 4A), in good agreement with our previous analysis^15^. Elimusertib induced dephosphorylation of ATR T1989 and its downstream target CHK1 S317 and a compensatory phosphorylation of ATM S1981 and its downstream targets CHK2 at S379 and S260. We observed a similar compensation at DNA-dependent protein kinase (DNAPK) sites S3205, S2612 and S2624. CHK1 phosphorylation at S286 was increased, likely reflecting CDK1/2 activity^26^. These responses were highly similar at high concentrations of ceralasertib (1 µM; 276 DP sites; Pearson’s *R* = 0.88), with the notable exception of ATR T1989, which responded more weakly to ceralasertib as compared to elimusertib (log2FC = - 0.42 versus log2FC = -1.74, respectively; Fig. 4A, C). In line with observations in NB cells (Fig. 3A), the DP response was largely absent at low concentrations (50 nM; 19 DP sites; Fig. 4A-B). SET S184 was one of the few sites that were dephosphorylated at low concentrations. Remarkably, this was also the strongest responding site at high concentrations (Table S1).

**Figure 4.**
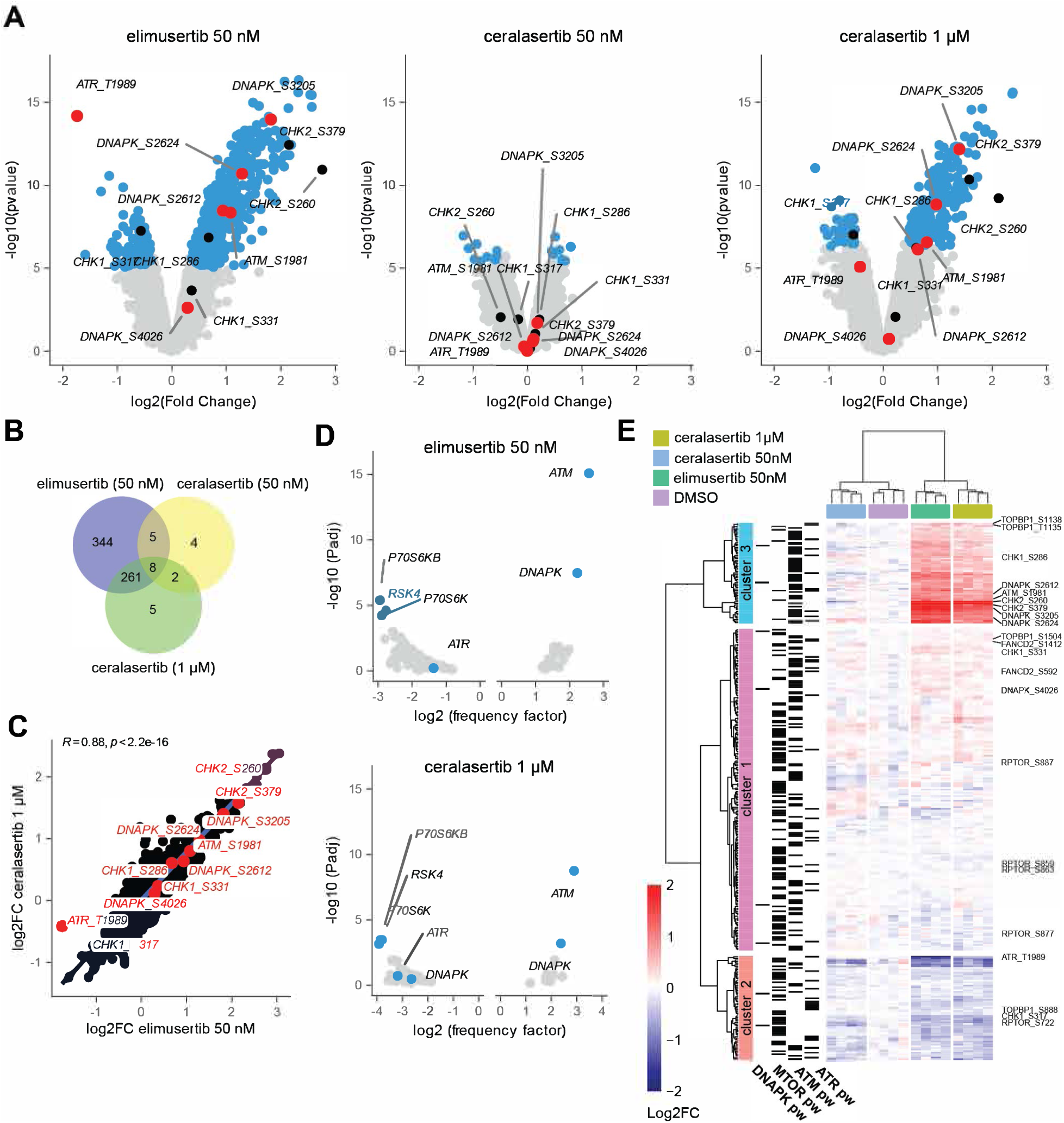
Comparison of the phosphoproteomic response to elimusertib and ceralasertib treatment of NB cell lines. CLB-BAR cells were synchronized by thymidine block and treated with either DMSO (control), elimusertib (50 nM) and ceralasertib (50 nM or 1 µM) for 6 h. **A.** Volcano plots showing differential phosphorylation (DP) between different treatments (as indicated) and control conditions. DP sites indicated in blue. ATR, ATM and DNAPK sites indicated and labelled in red. CHK sites indicated in black. **B.** Venn diagrams indicating number of DP sites for each condition. **C.** Correlation plot between DP log2FC values of different treatments as indicated. Sites in ATR, ATM, DNAPK and CHK1/2 indicated and labelled in red. Linear regression line in blue and Pearson correlation coefficient indicated on top. **D.** Volcano plot showing PK predictions for all hypophosphorylated (negative values) and hyperphosphorylated sites (positive values). ATR, ATM, DNAPK and strongest predicted PKs labelled and indicated in blue. **E.** Heatmap comparing log2FC between different samples as indicated. Hierarchical clustering performed using the Ward.2 method with dendrograms indicated. Genes active in ATR, ATM, DNAPK and mTOR pathways indicated on the left. See Table S1 for detailed results.

To identify the protein kinases mediating the DP response, we performed a motif enrichment analysis based on the recently published atlas of substrate specificities for the human serine/threonine kinome^27^. This analysis confirmed that ATM and DNAPK are the main drivers of the compensatory phosphorylation response. The strongest enrichments for the dephoshorylated motifs were observed for Ribosomal Protein S6 Kinases A6 (RSK4) and B1 (S6KB and S6K; Fig. 4D; Fig. S3A). While S6K is downstream of mTOR, the predicted downregulation of RSK4 is intriguing, as RSK4 is reported to be constitutively active in most cell types, in contrast to RSK1-3^28^.

We then focused on phosphorylation sites in different PIKK family members (ATM, ATR, DNAPK or mTOR). Using an unsupervised clustering approach, 3 obvious clusters were observed (Fig. 4E). Cluster 2 contained 65 sites that were mainly dephosphorylated upon elimusertib 50 nM and ceralasertib 1 µM treatment (mean log2FC = -0.37 and -0.38, respectively; P < 0.001 for both; Fig. S3B) and included ATR T1989, CHK1 S317 and RPTOR S722. This cluster was enriched for sites in proteins active in the ATR pathway (odds ratio = 2.8, *P* = 3.3e-03, Fisher’s exact test) as well as the mTOR pathway (odds ratio = 1.8, *P* = 0.037). Cluster 3 was enriched for proteins active in the ATM pathway (odds ratio = 5.26, *P* = 3.2e-08) and contained 63 phosphorylated sites, including ATM S1981, CHK2 S260 and S379, DNAPK S2612, S2624 and S3205 as well as CHK1 S286. Interestingly, while treatment with the low concentration of ceralasertib did not result in any response in the ATM-related cluster 3, a clear dephosphorylation (mean log2FC = -0.21; *P* = 1.1e-10) was still observed in the ATR-related cluster 2 (Fig. 4E).

In conclusion, our phosphoproteomic analysis suggests that ATR inhibition with either 50 nM elimusertib or 1 µM ceralasertib result in a highly similar reduction of ATR and mTOR signalling and a compensatory response that is driven by both ATM and DNAPK. This compensation was absent at low concentrations of ceralasertib.

### Use of elimusertib results in long term responses in ALK-driven NB mouse models

We next investigated the efficiency of these ATR inhibitors alone or in combination with lorlatinib in ALK-driven NB mouse models. For monotreatment, Alk/MYCN-driven GEMM tumours were treated with either 25 mg/kg elimusertib or ceralasertib twice daily for two 3-day periods over the 14-day protocol (Fig. 5A). For the combinatorial regimen the initial ATR inhibitor monotreatment (either elimusertib or ceralasertib) was followed by 10 mg/kg lorlatinib twice daily for 11 days, supplemented with the respective ATR inhibitor at days 8–10 (Fig. 5B). All mice tolerated the treatment regimen with no noticeable side effects. A similar tumour volume decrease was observed in response to both elimusertib and ceralasertib (Fig. 5C). In contrast to monotreatment, no tumours were detected at day 14 after treatment in any of the lorlatinib and elimusertib combinatorial treated GEMMs (Fig. 5C). However, in lorlatinib and ceralasertib combinatorial treated GEMMs we observed similar tumour volume decreases to that seen with elimusertib monotreatment (Fig. 5C). Taken together, these data indicate that the ALK inhibitor lorlatinib increases the anti-tumour efficacy of the ATR inhibitors elimusertib and ceralasertib, in our NB GEMMs.

**Figure 5.**
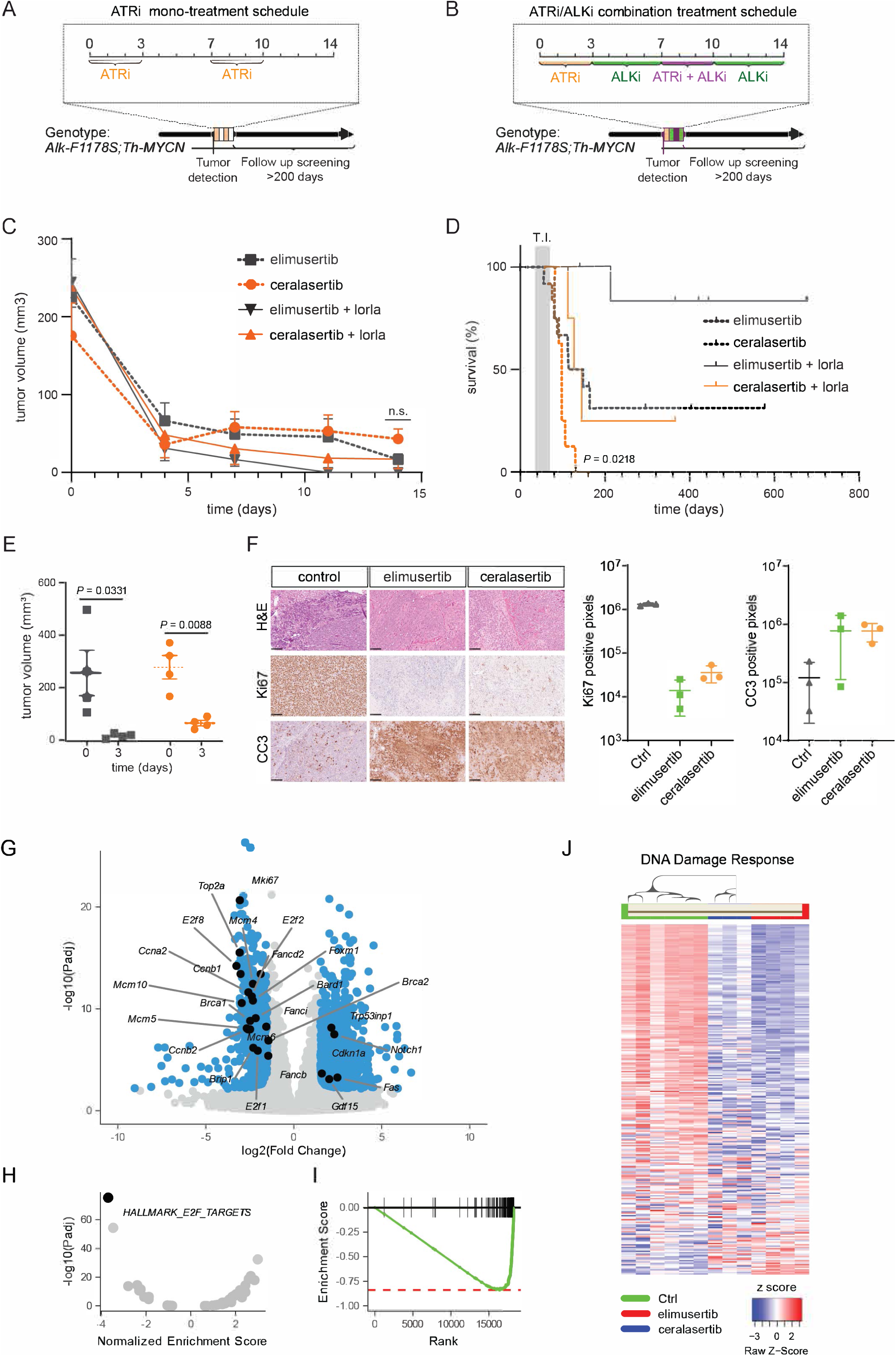
Elimusertib exhibits a more robust anti-tumour response in GEMMs compared to ceralasertib. **A.** Schematic overview of the monotreatment regimen for Alk/MYCN-driven GEMM tumours. Mice were treated with 25 mg/kg elimusertib or ceralasertib for 3 days. A subsequent 4 day pause from treatment was followed by an additional 3 days of ATRi. **B.** In the combination regimen the initial monotreatment with ATRi followed by 10 mg/kg lorlatinib for 11 days, supplemented with ATRi at days 8–10, orally. All treatments were given twice daily. **C.** Tumour volume was measured by ultrasound over 14 days. Data is presented as mean +/- SEM. **D.** Survival (from birth) of *Alk-F1178S;Th-MYCN* treated with elimusertib monotreatment (grey dashed, n=12) or ceralasertib (orange dashed, n=8) (*P*=0.0218) compared with elimusertib or ceralasertib and lorlatinib combination treatment (grey and orange solid respectively, n=7 and n=4) (*P*=0.025). Data for elimusertib and elimusertib + lorlatinib treatment cohorts are shown for comparison. Shadowed area shows normal tumour incidence range. Kaplan-Meier with logrank (Mantel-Cox test). **E.** Tumour volume, measured by ultrasound, after 3 days of either monotreatment with 25mg/kg elimusertib (grey squares, *P*=0.0331) or ceralasertib (orange circles *P*=0.0088). Paired one-tailed t-test, n=4. **F.** Immunohistochemistry of tumours in E, including an untreated control tumour of similar size, stained with H&E, Ki67 and Cleaved Caspase 3 (CC3, scale bar is 100 µm). Mean ± SD for Ki67 and CC3 positive pixels in tumours (n=3) for each treatment group. Each data point represents mean of three areas per tumour quantified with Ilastik. **G.** Volcano plots showing RNA-Seq based differential gene expression (DE) analysis between tumours from *Alk-F1178S;Th-MYCN* mice in untreated control conditions (n=6) and after 3 days of ceralasertib treatment (n=3). DE genes indicated in blue with genes discussed in main text indicated in black. **H-I.** Hallmark GSEA of DE analysis showing normalized enrichment scores and corresponding FDR values with labelling and running score plot (panel I) shown for E2F targets, the most strongly enriched gene set. **J.** Heatmap comparing z-score normalized gene expression counts between untreated control, ceralasertib treated and elimusertib treated *Alk-F1178S;Th-MYCN* mice for 273 different DDR genes as indicated. Genes ranked based on DE (most DE on top). Colour key shown at bottom right. Columns (tumour samples) hierarchical clustering according to the dendrogram shown on top.

Given this striking response to the 14-day elimusertib/lorlatinib regimen, we maintained all remaining mice over time without any therapeutic interventions, regularly monitoring tumour development with ultrasound. In addition, mice were subjected to recurrent follow-up by ultrasound, confirming complete tumour regression. All mice treated with ceralasertib alone relapsed within 30 days of treatment cessation. This is similar to the response rate observed by Roeschert and colleagues in a *Th-MYCN* driven NB mouse model^16^, although they employed 25-30 mg/kg ceralasertib, where all mice treated died during the 30 days treatment regime. In our hands, the ceralasertib/lorlatinib combination resulted in prolonged survival compared with ceralasertib alone, however, all mice except one relapsed within 50 days of treatment termination (Fig. 5D). In comparison, mice treated with elimusertib as a single agent exhibited increased survival when compared with either ceralasertib alone or ceralasertib/lorlatinib in combination, with three out of twelve mice remaining tumour free for up to 290 days after cessation of therapy (Fig. 5D). Further, and surprisingly, elimusertib monotreatment is superior to lorlatinib treatment alone in our Alk-driven NB models^15,19^.

Monotreatment with elimusertib had a significantly better survival rate as compared to ceralasertib (log-rank (Mantel-Cox) test, *P* = 0.0218; Fig. 5D). Remarkably, all but one *Alk-F1178S;Th-MYCN* mouse treated with the elimusertib/lorlatinib regimen have remained tumour free (for >350 days from treatment). The single relapsed mouse had a tumour in the lumbar region of the back 137 days after treatment cessation, at age 210 days. However, it is unclear whether this was a recurrence of the original tumour or a newly arising primary tumour. To further characterise the ATR inhibition response, tumours were treated with either elimusertib or ceralasertib for 3 days and sampled for histological and RNA-Seq analysis (Table S2). Treated tumours exhibited a reduced tumour volume (Fig. 5E), accompanied by reduced staining of the Ki67 proliferation marker and enhanced cleaved caspase 3 activity (CC3) when compared with vehicle controls (Fig. 5F). Similar to our previous analysis on elimusertib treated mice^15^, RNA-Seq analysis of *Alk-F1178S;Th-MYCN* tumours treated with ceralasertib displayed a downregulation of E2F targets and G2M checkpoint genes (Fig. 5G-J).

### Elimusertib treatment triggers a robust differentiation response in tumours

We next tested whether host immune cell involvement may assist in the rapid tumour loss on ATR inhibition. To investigate differences in immune cell infiltration, tumours from treated with elimusertib (25 mg/kg) or vehicle twice daily for 3 days were analysed by FACS with an immune cell panel. No obvious differences in immune cell infiltration of the tumour were observed, with similar immune cell levels from both innate and adaptive immune systems were found in the tumours, with intergroup variations greater than any treatment effect (Fig. S4). Subgroup analysis of CD4+ and CD8+ cells also failed to identify any significant changes between the two treatments or expression of the CD69 activation/residential memory marker. CD8 inhibition resulted in a slight decrease in tumour response to the 14 day combination treatment of elimusertib and lorlatinib in *Alk-F1178S;Th-MYCN* mice harbouring tumours (Fig. 6A). We also tested the effect of cGAS/STING inhibition, employing the H-151 inhibitor^29^. Notably, treatment with H-151 did not block the rapid tumour regression in mice receiving elimusertib and lorlatinib combination therapy, although it did result in a significant increase in relapse after treatment in comparison with elimusertib and lorlatinib (Fig. 6C). Thus, while immune cell responses appear to be important for the prolonged response observed in tumour bearing *Alk-F1178S;Th-MYCN* mice in response to combined ATR inhibition, neither CD8+ T cells nor the cGAS/STING response seem to be required for the rapid tumour response observed on ATR inhibition.

**Figure 6.**
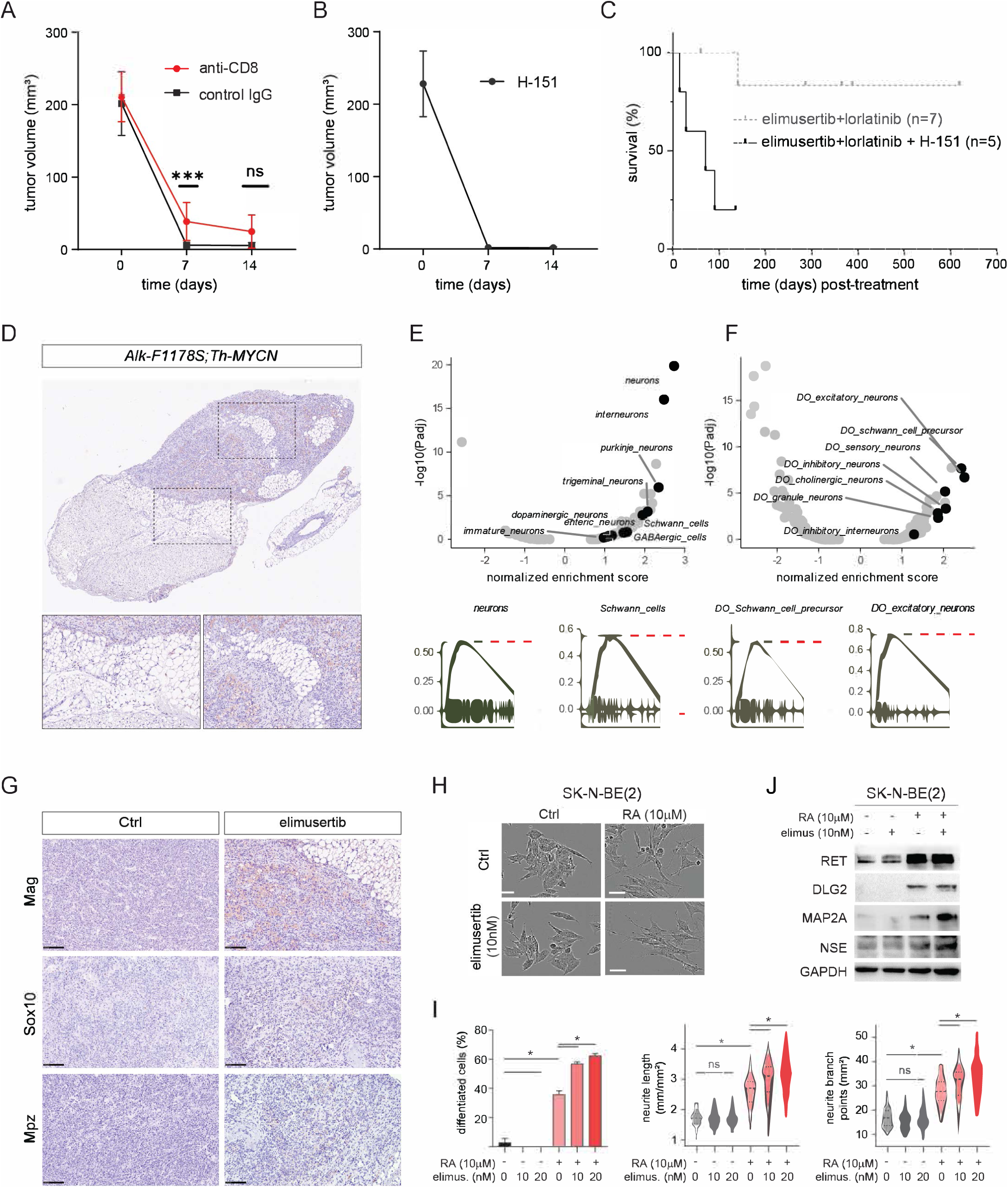
Investigation of immune cell dependency and differentiation upon ATRi treatment. **A.** Tumour volume in *Alk-F1178S;Th-MYCN* mice harbouring tumours treated with combination elimusertib and lorlatinib concurrent with injections of either anti-mouse CD8α (n=10 (day 0), n=10 (day 7), n=9 (day 14)) or IgG2b isotype control (n=5 (day 0), n=5 (day 7), n=4 (day 14)). Comparison of tumour volume differences between anti-mouse CD8α and IgG2b isotype controls at day 7 (*P*=0.0007) and day 14 (*P*=0.0755) determined by two-tailed Mann-Whitney test. **B.** *Alk-F1178S;Th-MYCN* mice harbouring tumours treated with H-151 (n=5) in addition to the 14 day combination treatment regime (elimusertib+lorlatinib). Tumour volume data presented as mean +/- SD. **C.** Kaplan-Meier survival curve (post-treatment) of *Alk-F1178S;Th-MYCN* mice harbouring tumours treated with H-151 (n=5) in addition to the 14 day combination treatment regime (elimusertib+lorlatinib). Gray dashed line shows survival curve for elimusertib and lorlatinib alone (Fig. 5D) for comparison. **D.** Representative H&E/Mag stained treated *Alk-F1178S;Th-MYCN* tumour, showing distinct histological features within the same tumour, inserts are magnifications of dashed squares. **E-F.** Volcano plots showing GSEA results using cell type signature genes from PanglaoDB **(E)** and MSigDB **(F)**. Enrichment was performed using DGE between NB tumours from *Alk-F1178S;Th-MYCN* mice in untreated control conditions (n=6) and after 3 days of elimusertib treatment (n=3). Neuronal and Schwann cell-related gene sets are indicated. Bottom plots showing running score plots for the most enriched neuronal and Schwann cell-related gene sets. **G.** Sections from control and elimusertib treated tumours were stained for Mag, Sox10 and Mpz by immunohistochemistry. Treated tumours exhibited Mag and Mpz positivity in areas with enlarged cells resembling neuronal or Schwann cells. Control tumours were negative. Sox10 positive nuclei were present in both control and treated tumours. **H.** Representative differentiation morphology images of SK-N-BE(2) cells upon DMSO, retinoic acid (RA)(10 μM) or/and elimusertib (10 nM) treatment for 24h. **I.** Bar graphs indicate percentage of differentiated cells, neurite length (mm/mm^2^), and neurite branch points (mm^2^) following RA or/and elimusertib treatment. **J.** Immunoblot for neuronal differentiation markers (RET, DLG2, MAP2, NSE) in SK-N-BE(2) cells treated with retinoic acid (RA) or/and elimusertib for 24 h.

Having been unable to identify a role for either the CD8+ T cells or the GAS/STING response in the rapid tumour regression response we conducted an in-depth histopathological examination of ATRi treated tumours. Careful inspection revealed tumour areas that exhibited characteristics of differentiation resembling neuronal or Schwann cell tissue (Fig. 6D), observed only in tumours treated with ATR inhibitors. Prompted by these findings, we searched for cellular differentiation signals in our tumour derived RNA-seq data, employing GSEA analysis using cell type signature gene sets derived from PanglaoDB and MSigDB. Remarkably, the strongest enrichments were found for neuronal gene sets and also Schwann cells (or their precursors) were significantly enriched in both datasets (Fig. 6E-F). Neuronal gene set enrichments were confirmed by focusing the analysis on gene sets derived from previous chemical and genetic perturbation experiments in mice, which also indicated strong depletion of genes upregulated in neuroendocrine lung carcinoma (Fig. S5).

Taken together, these data suggest that ATR inhibition triggers a differentiation response in mouse tumours. In agreement, a number of markers in the Schwann cell differentiation pathway were confirmed by immunohistochemistry. These included Mag and Mpz, which were expressed in *Alk-F1178S;Th-MYCN* tumours treated with elimusertib (Fig. 6G). We also noted increased Sox10 positivity on treatment with elimusertib, however, untreated control tumours also contained Sox10 positive cells in some areas. To test further the hypothesis that ATR inhibition triggers differentiation, we treated NB cells with low doses of ATR inhibitor in the presence or absence of retinoic acid (RA) and asked whether ATR inhibition was able to potentiate RA induced differentiation. While RA treatment induced neuronal differentiation in 36% of SK-N-BE(2) cells, addition of low dose elimusertib further induced neuronal differentiation to 63% in a dose-dependent manner (Fig. 6H-I). Neurite length and branch points were significantly increased by low dose elimusertib in the presence of RA, compared to RA-treated controls (Fig. 6I). Consistent with the neuronal differentiation phenotype, expression of established neuronal lineage differentiation markers, such as RET, DLG2, MAP2A and NSE, were further upregulated in by elimusertib in the presence of RA (Fig. 6J).

### Tumours treated with elimusertib exhibit differential DNA methylation

We next considered the possibility that ATRi-induced differentiation is due to an effect of the DDR on genome methylation. Epigenetic regulation, including DNA methylation, plays an important role in NB^30^, prompting us to examine changes in DNA methylation in *Alk-F1178S;Th-MYCN* tumours treated with elimuseritib for 72 h. Strikingly, immunohistochemical staining of ATRi-treated tumours for 5-methylcytosine (5-mC) revealed a considerable increase in DNA methylation, together with a Schwann cell-like phenotype (Fig. 7A). Subsequently, we conducted comprehensive whole-genome bisulfite sequencing (WGBS), and consistent with increased levels of 5-mC, identified a global augmentation in differentially methylated regions (DMRs) in ATR-inhibited tumours (Fig. S6A). This effect was evident across the entire genome, with elevated numbers of hypermethylated regions observed on each individual chromosome (Fig. 7B). The methylated regions had a peak length distribution close to 150bp (Fig. S6B), and quantification confirmed large widespread differences in methylation level between vehicle and ATRi-treated tumours (Fig. 7C). Specifically, a notable increase in methylated CpG islands and shores throughout the genome was observed, accompanied by intensified methylation within gene bodies, encompassing exons, introns, and untranslated regions (UTRs) in ATRi-treated tumours (Fig. 7D and Fig. S6C). Upon closer examination, the ratio between methylated CG (mCG) and unmethylated CG was larger within the gene body compared to the 2kb upstream and downstream regions of genes (Fig. S6D). Although ATR inhibition induced an overall increase in methylation levels, we also identified distinct genomic regions that exhibited higher levels of methylation, surpassing the DMR threshold of >0.5 (Fig. 7E). To gain insights into the cellular processes influenced by ATR inhibition through DNA methylation, we analysed highly hypermethylated gene bodies (DMR<0.5) identifying differences in DMR levels between elimuseritib and vehicle treated tumours (ΔDMR<0.2) (Fig. 7F). Remarkably, several among the ten most significant clusters encompassed methylated gene bodies associated with developmental processes, potentially illuminating the observed transition to a Schwann cell lineage in ATRi-treated tumours. Among interesting loci hypermethylated in response to ATRi, we noted *Dlx5*, which has previously been reported as a tumour initiation associated factor in NB. We also observed hypermethylation in components of the TGF-beta signaling and Ephrin pathway (e.g. *Tgfb1*, *Tgfbr3*, *Ephb3*, and *Ephb4*), which is interesting as previous research has shown that cross-talk between TGF-β and Ephrin pathways interferes with Schwann cell differentiation to drive peripheral nerve regeneration^31^. Additionally, we detected hypermethylation in *Fgfr2*, which is intriguing as inhibiting *Fgfr2* has shown promise in sensitizing *MYCN*-amplified NB to CHK1 inhibitors^32^. Hypomethylated regions were less frequent but still present following ATRi treatment (Fig. 7B) and cluster analysis revealed in total five significant clusters (Benjamini < 10^-6^) all involving cell adhesion, in keeping with previous reports of a link between adhesive properties of Schwann cells with levels of DNA methylation^33^. Collectively, our findings provide compelling evidence that ATR inhibition induces widespread hypermethylation in treated tumours, which is concurrent with the observed dedifferentiation process towards a Schwann cell-like phenotype. Further investigations are warranted to elucidate the precise roles of these genes in neuroendocrine cell differentiation and response to ATR inhibitors in NB.

**Figure 7.**
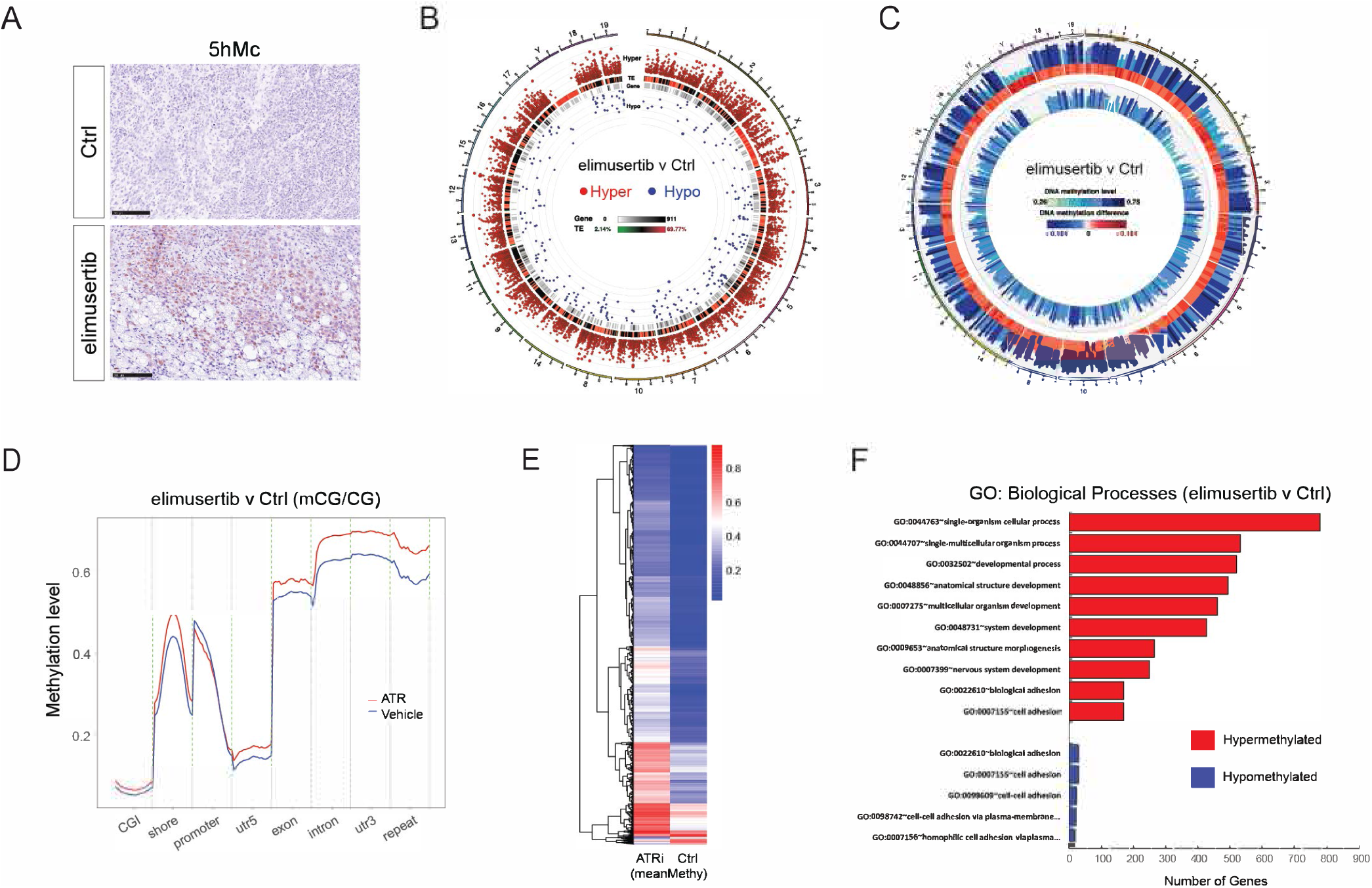
Elimusertib treated tumours exhibit a robust methylation response. **A**. Serial sections from elimusertib treated tumours were stained for H&E and 5-hydroxymethylcytosine (5-hmC) by immunohistochemistry. **B**. Circos plot showing hypo- and hypermethylated regions within the mouse genome between vehicle and elimusertib treated *Alk-F1178S;Th-MYCN* tumours. **C**. Circos plot depicting global differentially methylated regions (DMR) patterns within the mouse genome in vehicle and elimusertib treated *Alk-F1178S;Th-MYCN* tumours. **D**. Diagram showing DMR levels within functional genomic regions. **E**. Heatmap of DMR levels between vehicle and elimusertib treatment. **F**. Bar graph displaying significant hypo- and hypermethylated gene clusters in elimusertib treated *Alk-F1178S;Th-MYCN* tumours compared to vehicle. Vehicle treatment n=3, elimusertib treatment n=3 in all experiments. DMRs were sorted by areaStat including hypomethtlated (areaStat<0) and hypermethylated (areaStat>0) DMRs.

## Discussion

Management of high-risk NB presents a serious clinical challenge^8^. The identification of ALK mutations in NB, together with the synergistic effect of ALK with MYCN in driving aggressive and penetrant NB in both preclinical models and from clinical data, has raised hopes that ALK inhibition will be therapeutically beneficial in NB patients^4,34–37^. As a result, ALK mutation-positive NB patients are treated with ALK inhibitors, which typically elicit a strong response, and there have been reports of long-term responses^9,38–40^. However, patients treated with ALK TKIs also relapse with a complex landscape of potential resistance mutations^9,41,42^. Although the mechanisms behind ALK TKI resistance in relapsed NB patients are still not entirely understood, recent research suggests that a variety of mechanisms are involved, including the development of activating mutations in the RAS/MAPK pathway and ALK mutations themselves in ALK TKI-resistant tumours^42^. Therefore, discovering therapeutic approaches that enhance the efficacy of ALK TKI therapy in the clinical management of NB is a significant objective for the NB research community.

In recent years, ALK TKI combinatorial options have been identified in preclinical models, many with therapeutic potential for the clinic^15,43–46^. ATR is one such combination target, which has generated exceptional outcomes in preclinical ALK-driven models^15^. ATR inhibitors have been explored in NB, either as monotherapy, or in combination with PARP or Aurora-A inhibitors to block NB tumour and cell growth^16,17,47^. ATR inhibitors in clinical trial include M6620 and M4344 (Merck), ceralasertib (Astra Zeneca) as well as elimusertib (Bayer)^48,49^ and camonsertib (Repare Therapeutics/Roche)^50^ that are currently at various phases (www.clinicaltrials.gov). For example, ceralasertib in combination with anti-PD-L1 antibody has shown promising antitumour activity in phase II trials on metastatic melanoma, epithelial ovarian cancer and advanced gastric cancer patients ^51–53^.

The first reported combination of ATR inhibition with an ALK inhibitor employed a simple 14-day protocol that resulted in complete tumour resolution in ALK/MYCN driven NB GEMMs^15^. This can be compared to monotreatment with lorlatinib alone, which impairs tumour growth, but is unable to resolve tumours in these GEMMs^19^. Upon cessation of ALK inhibitor treatment, tumour growth resumes aggressively. Remarkably, in this study we show that ATRi monotreatment is sufficient to resolve ALK-driven tumours. Surprisingly, ATRi monotreatment displays better efficacy than the ALKi lorlatinib in an ALK/MYCN driven mouse model, while combining ATR and ALK inhibition resulted in a significantly increased long-term survival benefit. At the molecular level we have been able to identify the impact of ALK signalling on the DDR via phosphorylation of both ATR and CHK1. The ALK-dependent phosphorylation of CHK1-S280, rather than the ATR phosphorylation site (CHK1-S345), identifies an ALK-responsive site that has previously been reported to be downstream of both Akt and p90RSK^21,54^. Taken together, our findings here and in previous work^15,20^ suggest that ALK signalling in NB supports the robustness of the DDR. It is interesting to speculate whether the inhibition of ATR may also trigger feedback signalling resulting in increased dependency of the tumour on ALK signalling activity. Previous RNAi screening of kinases identified CHK1 as a therapeutic target in NB cell lines, also showing that high-risk NB exhibits increased CHK1 expression and activation^55^. Our identification of ALK-regulated phosphorylation of CHK1-S280 suggests that combination of ALK and ATR inhibition results in a more effective impairment of CHK1 function^56^.

In this study, we have compared two ATR inhibitors, elimusertib and ceralasertib, and show that both drugs are effective in resolving ALK-driven tumours in our GEMMs. However, tumour bearing animals treated with ceralasertib relapse significantly faster than those treated with elimusertib. Our phosphoproteomics analyses also suggest that elimusertib is more potent in NB cells, with 1 µM ceralasertib resulting in similar, but not identical, phosphoproteomic profiles to 50 nM elimusertib. Both drugs led to compensation of ATR inhibition by upregulation of ATM and DNAPK activity. We have previously noted the compensatory activation of ATM on inhibition of ATR^15^. In the present study we have employed the recently reported atlas of substrate specificities for the human serine/threonine kinome^27^, which identified increased DNAPK activity in response to ATR inhibitor treatment. This is further supported by our observation of increased phosphorylation on DNAPK-S3205, which has been reported as both a DNAPK autophosphorylation site^57^, and a target of ATM^58^.

A key finding in this work is the striking differentiation of tumours in response to ATR inhibition. Induction of differentiation in NB is considered a therapeutically exploitable option, with treatments such as retinoic acid employed in the clinic to promote NB differentiation^59–61^. Our initial ATR study focused on the impressive ability of ATR inhibition to induce cell death in a wide range of NB cell lines as well as in tumours in ALK-driven GEMMs. The rapid reduction in tumour size led us to investigate a role for the immune system in this process. However, although inhibition of T cells with anti-CD8 antibodies had some effect, it failed to block tumour shrinkage in response to ATR inhibition. Likely, DC-mediated activation of the T-cell compartment plays an important role in the long-lasting response observed in these mice, an aspect that will be important to investigate in future studies. Interestingly, a recent study has reported that ATRi treatment cessation can potentiate T cell responses, suggesting that our 14 day schedule may be beneficial ^62^. Given that targeting of the DDR leads to activation of the cGAS-STING pathway, we also investigated whether blocking this pathway with the H-151 inhibitor that blocks activation-induced palmitoylation of STING^29^, would affect tumour shrinkage in response to ATR inhibition. Again, while we show that activation of cGAS-STING appears to be important for the long-term prevention of relapse, we could not obtain any strong evidence that this pathway plays an important role in the rapid tumour response.

In the current era of high-tech data analyses, it is easy to overlook simple methodology. Routine histological analysis of tumours treated with ATR inhibitors revealed large regions of tumour that resembled adipose or neural tissue. This was confirmed histologically and through reanalysis of RNA-seq data from treated tumours. RNA-seq analyses identified a strong enrichment of neuronal and Schwann cell gene sets. Interestingly, earlier work has shown that deletion of *Atr* in adult mice leads to depletion of stem and progenitor cells^63^. This loss of stem and progenitor cells has been suggested to arise due to elevated apoptosis or senescence, however, no differences could be observed between *Atr*-deficient and control tissues, raising the possibility that deletion of *Atr* triggered aberrant, ectopic differentiation, leading to the loss of proliferating stem and progenitor cells. This hypothesis is supported by a recent report in the fruit fly, *Drosophila melanogaster*, that identified a critical role for *Drosophila* ATR (*mei-41*) in preventing neuroblast differentiation in response to irradiation stress^64^.

Interestingly, inhibition of the ATR and ATM DDR kinases has been linked with differentiation in acute myeloid leukemia cells carrying MLL fusion proteins, where treatment with ATR inhibitors led to terminal differentiation and loss of leukemic blasts^65,66^. We show here that low dose elimusertib synergises with retinoic acid, which is currently employed clinically to promote NB differentiation^60,61^. This observation suggests that ATR inhibition may increase the potential of differentiation therapy in NB and highlights the need to elucidate the mechanisms underlying the dramatic ATR inhibition-induced differentiation response in our NB tumour models. Ultimately, ATR inhibition may promote multiple important tumour effects, by promoting NB differentiation as well impairing the DDR leading to mitotic catastrophe. The effect of ATR inhibition on the tumour epigenome we observed *in vivo* is supported by many studies linking the DDR and genome methylation. While detailed investigation is beyond the scope of this study, we note differential methylation of many loci involved in neuronal differentiation that will be interesting to examine in future work.

Our findings show that this ATR-ALK 14-day combination treatment is highly effective in ALK-driven GEMM models of NB. However, several questions remain. For example, is this treatment useful for non-ALK driven NB? *Th-MYCN* tumours exhibit high levels of replication stress, which suggests they would be sensitive to ATR inhibition, and indeed treatment with ceralasertib has already been reported in preclinical models, although all treated tumour bearing mice died during the treatment period^16^. To our knowledge, there are no reported clinical NB cases in which ATR inhibitors have been employed, and the question of which patients may benefit most is important to consider. High-risk NB patients exhibiting *MYCN*-amplification, *ALK*-activating mutations and overexpression/amplification, combined *MYCN*/*ALK* perturbations and 2p-gain, are all NB categories that potentially exhibit high levels of oncogene-induced replicative stress and be sensitive to ATR inhibition. An additional high-risk NB category of interest are the 11q-deletion group, where many genes involved in the DDR, such as *ATM*, *CHEK1*, *H2AFX* and *MRE11*, are lost. In our previous study, we observed that all NB cell lines tested, irrespective of *ALK*, 11q or *MYCN* amplification status, were sensitive to inhibition of ATR with elimusertib. In future work it will be interesting to focus on the potential of ATR inhibition on these different genetic categories. It will also be important to consider means of identifying NB cases that may respond to ATR inhibition. In this respect, a recent study has identified nuclear pCHK1 as a potential biomarker of ATR sensitivity to ATR inhibition^67^.

NB tumours harbouring activating RAS mutations such as *NRASQ61\** or *KRASQ61\**^5,68^ would also be expected to exhibit high levels of replication stress, and 50 nM elimusertib is highly effective in preventing growth of SK-N-AS NB cells^15^, making it of interest to test whether this ATRi/ALKi combination, or perhaps an ATRi/MEKi combination, would be effective in a *RAS* mutant NB setting. This is of particular interest considering recent findings that have identified potential *RAS* mutant resistance mutations in lorlatinib treated NB patients^9^.

Ultimately, this work identifies ALKi/ATRi combination therapy as an effective treatment in ALK-driven NB GEMMs. We further show that ALK signalling primes the DDR through phosphorylation of both ATR and CHK1, providing a molecular mechanism underlying the efficacy of this combination. The strong neuronal and Schwann cell signatures observed in tumours treated with ATR inhibitors, together with increased DNA methylation levels, suggest that ATR inhibition exerts a robust differentiation response that underlies tumour regression. Taken together, these results strongly motivate the continued exploration of ATR/ALK inhibition as a potentially therapeutically effective approach in NB.

## Methods

### Antibodies and inhibitors

Primary antibodies against ATR (#13934, 1:1000), pATR (S428, #2853, 1:1000), ATM (#2873, 1:1000), pATM (#13050, 1:1000), CHK1(#2345, 1:1000), pCHK1(S345,#2348, 1:1000), pCHK1(S280,#2347, 1:1000), pAKT (S473,#4060, 1:4000), DLG2 (#19046), pERK1/2 (Y204/T202, #4377, 1;2000), pALK (Y1278, #6941, 1:1000), β-Actin (#4970, 1:10000), AKT (#9272), FOXM1 (#20459, 1:1000), pFOXM1 (T600, #14655, WB 1:1000, IF 1:400, IHC 1:1000), Ki67 (#12202, 1:800), MAP2A (#8707), RET (#3220)α-tubulin (#2125, 1:10000), cleaved caspase-3 (#9661, 1:200), p21 (#2947, 1:50), survivin (#2808, 1:400), phospho-histone H3 (9701, 1:200), GAPDH (#5174, 1:20000), p53 (#2527, 1:1000) and PARP (#9542, 1:1000) and were obtained from Cell Signalling Technology. Monoclonal antibody 135 (anti-ALK, 1:100) was produced in-house against the extracellular domain of ALK^69^. Antibodies against SOX10 (ab180862, 1:400), 5-hydroxymethylcytosine (5-hmC) (ab214728, 1:15000) NSE (#ab53025), and Myelin Protein Zero (MPZ) (ab183868, 1:200), were obtained from Abcam. Polyclonal antibodies against MAG/Siglec-4a (NBP2.17201, 1:250) were obtained from Novus. Horseradish peroxidase– conjugated secondary antibodies, goat anti-mouse immunoglobulin G (IgG) (# 32230), and goat anti-rabbit IgG (# 32260, 1:5000) were purchased from Thermo Fisher Scientific. Lorlatinib (RLor-10-21) was purchased from Reagency (Melbourne, Australia), elimusertib (HY-101566) and ceralasertib (HY-19323) were purchased from MedChemTronica (Sollentuna, Sweden). Etoposide (S1225) was purchased from Selleckchem (Houston, USA) and LY2603618 (T6084) was from TargetMol Chemicals Inc. (Boston, MA, USA). InVivoPlus anti-mouse CD8α (#BP0061) and InVivoPlus rat IgG2b isotype control, anti-keyhole limpet hemocyanin (#BP0090) were purchased from BioXcell (Lebanon, USA). H-151 (T5674) was purchased from TargetMol (Wellesley Hills, U.S.A).

### Cell culture

CLB-BAR, CLB-GE, CLB-GAR, SK-N-BE(2), IMR32 and NB-1 NB cell lines were employed in this study. CLB-BAR (gain of function, Δexon4–11 truncated ALK), CLB-GE (gain of function, *ALK-F1174V* mutation), and CLB-GAR (gain of function, *ALK-R1275Q* mutation, 11q deletion) were obtained from The Centre Leon Bernard, France under MTA. These cells were cultured on collagen coated plates. IMR-32 (wild-type ALK, ligand-dependent activation) were purchased from ATCC. All cell lines were cultured in complete media, RPMI 1640 supplemented with 10% foetal bovine serum (FBS) with 1% penicillin/streptomycin at 37°C and 5% CO_2_. For ALKAL2 stimulation CLB-BAR and NB1 were used. Cells were seeded in 12-well plates and treated with inhibitors (2 h) or ALKAL2 ligand (0.5 h) alone or in combination. The ALK inhibitor lorlatinib (20 nM), ATR inhibitor elimusertib (100 nM), and the CHK1 inhibitor LY2603618 (1 uM) were used. For combination treatment, the cells were treated with inhibitors for 1.5 h prior to stimulation with ALKAL2 for half an hour. Cells were directly lysed in 1x SDS sample buffer and resolved on 8% SDS-PAGE. Primary antibodies, including pALK Y1278, ALK, pCHK1 S280, pCHK1 S345, pAKT S473, pERK1/2 and β-tubulin, were used to detect protein phosphorylation and total protein levels. For lorlatinib inhibition, CLB-BAR, CLB-GE, and CLB-GAR cells were seeded into 6-well plates and treated with lorlatinib (30 nM) for 0, 1, 6 and 24 h. Primary antibodies against pALK Y1278, ALK, pCHK1 S280, pCHK1 S345, CHK1, pH2A.X, Actin, and GAPDH were used. For etoposide treatment, CLB-BAR, CLB-GE, and CLB-GAR cells were seeded in 6-well plates and treated with lorlatinib (30nM) or etoposide (500nM), either alone or in combination for 24 h. DNA damage was monitored by immunoblotting for p-H2A.X S139. GAPDH was employed as a loading control. For synchronisation, cells were seeded on pre-coated collagen plates at 37°C and 5% CO_2_ overnight. Thymidine was added to the final concentration of 2 mM for 24 h. After 24 h cells were washed with fresh media prior treatment with either DMSO or ATR inhibitors.

### Subcellular fractionation

The Subcellular Protein Fractionation Kit for Cultured Cells (#78840, Thermofisher) was used following the manufacturer’s protocol. In brief, CLB-BAR cells in 10 cm dishes were treated with either DMSO control, lorlatinib or ALKAL2 respectively and then harvested for fractionation. Concentration of cytoplasmic and nuclear extracts were determined with a Pierce™ BCA Protein Assay Kit and 10 ug analysed on 8% SDS-PAGE. pCHK1 S280 antibody was used to detect the phosphorylation of CHK1, while pan CHK1 was used to detect total protein levels. Nuclear protein marker Lamin A and cytoplasmic protein marker β-tubulin were used as loading and fractionation controls.

### Immunoprecipitation and immunoblotting

For immunoblotting, protein lysate and immunoprecipitation samples were separated on 7.5% bis-acryl-tris gels, transferred to polyvinylidene difluoride membranes (Millipore), blocked in 3% bovine serum albumin (BSA) (phosphoprotein blots) and immunoblotted with primary antibodies as indicated overnight at 4°C. Secondary antibodies were diluted 1:10000 and incubated at room temperature for 1 hour. Enhanced chemiluminescence substrates were used for detection (GE Healthcare), and membranes were scanned using LI-COR Odyssey instrumentation.

### Fluorescence microscopy

For pFOXM1 T600 imaging, cells were fixed with 4% PFA/PBS for 10 min, and permeabilized for 10 min with ice-cold methanol, blocked in 1% BSA/PBS for 30 min at RT. The primary antibodies were diluted in 1% BSA/PBS and incubated overnight at 4°C. Cells were washed 3x with PBS. Secondary antibodies (diluted 1:1000) and DAPI (5 μg/mL) were diluted in 1% BSA and incubated for 1 h at RT. Cells were washed 3x with PBS. A Zeiss LSM 800 confocal microscope and ZEN 3.5 Blue edition software were used to acquire images.

### IC_50_ determination

CLB-BAR and CLB-GE cells were seeded on 48-well plates. Next day, cells were treated with increasing concentration of elimusertib and ceralasertib ATR inhibitors. Cellular viability was assessed at 72 h with resazurin assay, with cells incubated with 44 µM resazurin (Sigma Aldrich) for 2- 3 h, followed by fluorescence (λex/em = 490/595 nm) quantification in a TEKAN microplate reader.

### Preparation of samples for phosphoproteomics analyses

Phosphoproteomics was performed on cells synchronised by thymidine block (2mM thymidine, 24 h). Briefly, 10.5 × 10^6^ cells (CLB-BAR) were seeded in FBS-free RPMI 1640 cell culture media in 15 cm^3^ dishes. One 15 cm^3^ dish of cells was used for each replicate, and experiments were performed in triplicate. After 24 h cells were either (i) subjected to thymidine block (2 mM thymidine, 24 h), or (ii) subjected to thymidine block (2 mM thymidine, 24 h), then washed twice with FBS-free RPMI, then released for 6 h in the presence of elimusertib (50 nM), ceralasertib (50 nM and 1 µM) or DMSO control. Cells were harvested by washing with PBS, briefly trypsinising and then neutralizing with non-FBS containing media. Aliquots were removed to perform immunoblotting validation and the remaining cells isolated by centrifugation at 200 x g, for 6 min. Pellets were resuspended in 1 ml of ice-cold PBS, and cell washing was repeated 5 times. Cell pellets were stored at -80°C for subsequent analysis.

Cells were lysed in 2M Guanidinium(Gua)-HCl, 0.1M ammonium bicarbonate, phosphatase inhibitors (Sigma P5726&P0044) by sonication (Bioruptor, 10 cycles, 30 seconds on/off, Diagenode, Belgium) and proteins were digested as described previously^70^. Shortly, 200 ug of protein per sample were reduced with 5 mM TCEP for 10 min at 95 °C and alkylated with 10 mM chloroacetamide for 30 min at 37 °C. After diluting samples with 100 mM ammonium bicarbonate buffer to a final Gua-HCl concentration of 0.5 M, proteins were digested by incubation with sequencing-grade modified trypsin (1/50, w/w; Promega, Madison, Wisconsin) for 12 h at 37°C. After acidification using 5% TFA, peptides were desalted using C18 reverse-phase spin columns (Macrospin, Harvard Apparatus) according to the manufacturer’s instructions, dried under vacuum and stored at -20°C until further use. Peptide samples were enriched for phosphorylated peptides using Fe(III)-IMAC cartridges on an AssayMAP Bravo platform as recently described^71^.

### Sample preparation for proteomic analysis using Tandem Mass Tags

Samples were homogenized on a FastPrep®-24 instrument (MP Biomedicals, OH, USA) for 5 repeated 40 s cycles at 6.5 m/s in lysis buffer containing 2 % sodium dodecyl sulfate (SDS), 50 mM triethylammonium bicarbonate (TEAB). Lysed samples were centrifuged at 8000 xg for 20 min and supernatants transferred to clean tubes. Protein concentrations were determined using Pierce™ BCA Protein Assay Kit (Thermo Fisher Scientific) and the Benchmark™ Plus microplate reader (Bio-Rad Laboratories, Hercules, CA, USA) with bovine serum albumin (BSA) solutions as standards. Aliquots containing 600 µg of protein from each sample were incubated at room temperature for 60 min in lysis buffer with 100 mM DL-dithiothreitol (DTT). Reduced samples were split into two aliquots and processed using the modified filter-aided sample preparation (FASP) method^72^. Briefly, samples were transferred to 30 kDa Microcon Centrifugal Filter Units (cat no. MRCF0R030, Merck), washed repeatedly with 8 M urea and once with digestion buffer prior (0.5 % sodium deoxycholate (SDC) in 50 mM TEAB) to alkylation with 10 mM methyl methanethiosulfonate in digestion buffer for 30 min. Digestion was performed in digestion buffer by addition of Pierce MS grade Trypsin (Thermo Fisher Scientific), in an enzyme to protein ratio of 1:100 at 37°C overnight. An additional portion of trypsin was added and incubated for 4 h. Peptides were collected by centrifugation and the two aliquots per sample were pooled again. Isobaric labeling was performed using Tandem Mass Tag (TMTpro 16plex) reagents (Thermo Fisher Scientific). Labelled samples were combined into one pooled TMT set, concentrated using vacuum centrifugation, and SDC was removed by acidification with 10 % TFA and subsequent centrifugation. Peptide samples were purified using Pierce peptide desalting spin columns (Thermo Fisher Scientific) according to the manufacturer’s instructions.

The 9,600 µg peptides in the pooled sample were split into 540 µg for whole proteomics and two equal aliquots for phosphopeptide enrichment. The first aliquot was subjected to High-Select TiO2 Phosphopeptide Enrichment Kit, whereas the second aliquot was enriched by High-Select Fe-NTA Enrichment Kit (both Thermo Fisher Scientific) in parallel. Eluted phosphopeptide samples from both enrichments were pooled and fractionated into 14 fractions by increasing ACN concentration from 7 % to 50 % using the Pierce High pH Reversed-Phase Peptide Fractionation Kit (Thermo Fisher Scientific). Peptide separations of the whole proteome sample were performed using basic reversed-phase chromatography (bRP-LC) with a Dionex Ultimate 3000 UPLC system (Thermo Fisher Scientific) on a reversed-phase XBridge BEH C18 column (3.5 μm, 3.0×150 mm, Waters Corporation); using a gradient from 3% to 55 % solvent B over 45 min. Solvent A was 10 mM ammonium formate buffer at pH 10.00 and solvent B was 90 % acetonitrile, 10 % 10 mM ammonium formate at pH 10.00. The initial 50 fractions were pooled into 20 fractions, dried and reconstituted in 3 % acetonitrile, 0.2 % formic acid.

### nLC-MS analysis

The fractions of the whole proteome sample were analysed on an Orbitrap Lumos™ Tribrid™ mass spectrometer interfaced with Easy-nLC1200 liquid chromatography system (Thermo Fisher Scientific). Peptides were trapped on an Acclaim Pepmap 100 C18 trap column (100 μm x 2 cm, particle size 5 μm, Thermo Fisher Scientific) and separated on an in-house packed analytical column (75 μm x 35 cm, particle size 3 μm, Reprosil-Pur C18, Dr. Maisch). The whole proteomics fractions were separated using a gradient from 5 % to 35 % B over 77 min at a flow of 300 nL/min. Solvent A was 0.2 % formic acid and solvent B was 80 % acetonitrile, 0.2 % formic acid.

MS scans for whole proteomics were performed at a resolution of 120,000, and an m/z range of 375-1500. MS/MS analysis was performed in a data-dependent, with top speed cycle of 3 s for the most intense doubly or multiply charged precursor ions. Precursor ions were isolated in the quadrupole with an isolation window of 0.7 m/z, with dynamic exclusion set to 10 ppm and duration of 45 s. Isolated precursor ions were subjected to collision induced dissociation (CID) at a collision energy of 30 % with a maximum injection time of 60 ms. Produced MS2 fragment ions were detected in the ion trap followed by multinotch (simultaneous) isolation of the top 10 most abundant fragment ions for further fragmentation (MS3) by higher-energy collision dissociation (HCD) at 55 % and detection in the Orbitrap at a resolution of 50,000, m/z range of 100-500. Phosphoproteomics samples were separated using a linear gradient from 7 % to 35 % B over 47 min. Analysis was performed as above, besides that the MS2 analyses were split depending on mass and charge: MS2 scans of precursors 375-1375 m/z were assigned priority 1 (max. inj. time 200 ms). With 375-900 m/z and 3-6 charges, the isolation window of 1 was used, precursors 900-1375 m/z and charges 2-6, the isolation window was set on 1.2. Priority 2 were MS2 scans of 600-900 m/z and charge state 2, which had isolation window of 1 and 150 ms injection time. MS2 scans of 500-600 m/z and charge 2 precursors were assigned priority 3 and collected at isolation window of 1 and 100 ms injection time. Isolated precursors for this phosphoproteome were fragmented by higher-energy collision dissociation (HCD) using a collision energy of 38 % each. MS2 spectra were detected in the Orbitrap with the fixed first m/z of 100.

### Data analysis of Tandem Mass Tags phosphoproteomics

The data files of the fractions from each set were merged for identification and relative quantification using Proteome Discoverer version 2.4 (Thermo Fisher Scientific). Identification was performed using Mascot version 2.5.1 (Matrix Science) as a search engine by matching against the Homo sapiens database from Swiss-Prot (from January 2022). The precursor mass tolerance was set to 5 ppm and fragment mass tolerance to 0.6 Da (total proteome) or 30 mmu (phosphoproteome). Tryptic peptides were accepted with zero missed cleavages for whole proteomics, and variable modifications of methionine oxidation; fixed cysteine alkylation, and TMTpro-label modifications of N-terminus and lysine were selected for both. For the phosphoproteomics samples, variable modification of serine, threonine, or tyrosine phosphorylation was added. Phosphopeptides were identified in three subsequent steps: one missed cleavage, two missed cleavages, one missed cleavage and deamidation of asparagine or glutamine with 2 ppm precursor mass tolerance. Percolator was used for the validation of identified proteins.

For quantification, TMTpro reporter ions were identified with 3 mmu mass tolerance in the MS2 HCD spectra for the phosphoproteome or in the MS3 HCD spectra for the total proteome. The S/N values of the TMTpro reporters for each sample were normalized within Proteome Discoverer 2.4 on the total peptide amount. Only unique peptides, and those with a S/N ratio above 10, were considered for the protein quantification (whole proteome with minimum SPS match of 40 %). Peptides were filtered for high confidence and resulting identified proteins for medium confidence.

### Phosphoproteomics DIA LC-MSMS analysis

Phospho-enriched peptides were resuspended in 0.1% aqueous formic acid and 0.2 ug of peptides subjected to LC–MS/MS analysis using an Orbitrap Exploris 480 Mass Spectrometer fitted with an Vanquish Neo (both Thermo Fisher Scientific) and a custom-made column heater set to 60°C. Peptides were resolved using a RP-HPLC column (75μm × 30cm) packed in-house with C18 resin (ReproSil-Pur C18–AQ, 1.9 μm resin; Dr. Maisch GmbH) at a flow rate of 0.2 μLmin-1. Separation of peptides was achieved using the following gradient: 4% Buffer B to 10% Buffer B in 5 min, 10% Buffer B to 35% Buffer B in 45 min, 35% Buffer B to 50% Buffer in 10 min. Buffer A was 0.1% formic acid in water and buffer B was 80% acetonitrile, 0.1% formic acid in water. The mass spectrometer was operated in DIA acquisition mode with a total cycle time not exceeding approximately 3 s. For MS1, the following parameters were set: Resolution: 120,000 FWHM (at 200 m/z), Scan Range: 350-1400 m/z, Injection time: 25 ms, Normalized AGC Target: 300%. MS2 (SWATH) scans were acquired using the following parameters: Isolation Window: 8 m/z, HCD Collision Energy (normalized): 28%, Normalized AGC target: 1000%, Resolution: 15,000 FWHM (at 200 m/z), Precursor Mass Range: 400 – 1200 m/z, Max. Fill Time: 22 ms, DataType: Centroid. In total 100 DIA (MS2) mass windows per MS cycle followed by the one MS1 scan.

### Data analysis of phosphoproteomics DIA-MS data

The acquired raw files were searched using SpectroNaut (v17.1, PTM workflow, default settings) against a human database (consisting of 20360 protein sequences downloaded from Uniprot on 20220222) and 392 commonly observed contaminants using the following search criteria: full tryptic specificity was required (cleavage after lysine or arginine residues, unless followed by proline); 3 missed cleavages were allowed; carbamidomethylation (C) was set as fixed modification; oxidation (M), N-acetlyation (N-term) and phosphorylation (STY) were applied as variable modifications. The normalized, phosphosite centric and statstically analysed quantiative results were exported as a csv-table from the SpectroNaut software using the candidate list export option without any filters applied.

### Proteomics differential expression analysis

Differential protein expression was determined using the R Differential Enrichment of Proteomics data (DEP) package (default settings)^73^. Proteins and phosphosites were considered differentially expressed/phosphorylated when the absolute log2 fold change values were above 0.3 at FDR-adjusted *P* values below 0.05, respectively, considering a hyperbolic threshold as described earlier^15^.

### Protein kinase enrichment analysis

Protein kinase enrichment analysis was performed using the atlas of substrate specificities for the human serine/threonine kinome as suggested previously^27^. Briefly, for each phosphorylation site, the top 1% predicted protein kinases were derived from atlas of substrate specificities. A contingency table was then created comparing the proportion of differentially phosphorylated sites (hypo- or hyperphosphorylated, as defined higher) for each protein kinase. Statistical significance was evaluated using a one-sided Fisher’s exact test followed by multiple testing correction using the Benjamini-Hochberg method. Predicted protein kinases were plotted in a Volcano plot using the relative risk scores (log2 scale, x-axis; referred to as „frequency factors”) and adjusted *P* values (log10 scale, y-axis). Relative risk scores derived from hypophosphorylated sites were given an arbitrary negative relative risk score and only PKs with an adjusted *P* value below 0.1 were plotted for visualization purposes.

### Transgenic mouse model survival

*Alk-F1178SKI/0;TH-MYCNTg/0* (n = 31, mixed sex) on a 129×1/SvJ background (JAX stock #000691) were screened by ultrasound, 2–3 times a week from weaning. Tumours were followed until a size of 150-350mm^3^ in volume, at which a 3D scan of the tumour was made and treatment started (Day 0). Screening and 3D image acquisition was done utilizing Vevo 3100 imaging systems (FUJIFILM VisualSonics, Toronto, Canada). The VisualSonics MX550D (25–55MHz, 40 μm axial resolution) linear array transducer was used for all image acquisition. For 3D scanning, the probe was attached to a step motor, animals were anaesthetised with isoflurane and respiration rate and ECG were monitored during the procedure. Images were analysed in VevoLab (Fujifilm VisualSonics, Toronto, Canada). Combination treatment consisted of 3 days ATR inhibitor (elimusertib or ceralasertib diluted in 2% DMSO + 30% PEG 300) 25 mg/kg P.O. twice daily, 4 days lorlatinib 10 mg/kg (diluted in 2% DMSO + 30% PEG 300) twice daily, 3 days of both inhibitors as previously and 4 days of monotreatment with lorlatinib, for a total of 14 days. Tumour volume calculations, emplying the formula for an ellipsoid 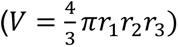 or the open multi-slice method in VevoLab, were performed on Day 0, Day 4, Day 7, Day 11, Day 14 and at sacrifice by meusuring the diameter in three dimensions from data obtained by ultrasound or caliper measurement. At the end of treatment mice were followed with ultrasound or until tumour could be clearly observed without palpitation, deteriorated health of the mice or sudden death. Humane endpoints were a tumour size exceeding in average diameter 20mm or if the animal showed symptoms due to tumour burden, mice that met these criteria were sacrificed. All experimental procedures and protocols were performed in accordance with the Regional Animal Ethics Committee approval, Jordbruksverket (Id.nr. 003225, Dnr 5.8.18-11644/2020, Id.nr. 004793, Dnr 5.8.18-02319/2023).

### Tumour preparation for downstream analyses

*Alk-F1178SKI/0;TH-MYCNTg/0* animals with tumours between 160-910 mm^3^, were treated for 3 days with ATR inhibitor (elimusertib or ceralasertib, 25 mg/kg) or vehicle twice per day orally. Mice were then sacrificed and tumours harvested, tumour material from each tumour were preserved in RNAlater (AM7024, Fisher Scientific, Göteborg, Sweden) for sequencing, 10% Neutral Buffered Formalin (HT5011, Merck KGaA, Darmstadt, Germany) for histology or snap frozen in liquid nitrogen for whole-genome bisulfite sequencing.

### Histological analysis of tumour sections

Tumour tissue were fixed in 10% neutral buffered formalin (HT501128, Merck KGaA, Darmstadt, Germany), embedded in paraffin, and sections of 5 μm were prepared by using microtome. Paraffin sections were stained with hematoxylin and eosin (H&E; Histolab Products) for morphological analysis. Proliferation and apoptosis were analysed by immunohistochemistry of Ki67, and cleaved caspase 3. Schwann cell markers were analysed by immunohistochemistry against Sox10, Mpz and Mag, and DNA methylation by antibodies against 5-hmC. Primary antibodies were detected with SignalStain® Boost IHC Detection Reagent (HRP, Rabbit) (#8114, Cell Signalling Technology), developed with SignalStain® DAB Substrate Kit (#8059, Cell Signalling Technology) and counterstained with hematoxylin (#01820, Histolab Products AB). Sections from three separate tumours in each treatment group (elimusertib, ceralasertib and vehicle) were scanned after staining as previously described with a Hamamatsu NanoZoomer-SQ Digital slide scanner. In short, three areas (of 1920 x 1216 pixels) of each section were obtained utilizing the NanoZoomer Digital Pathology viewer 2 and subsequently analysed and quantified for positive pixels with the interactive machine learning software Ilastik^74^.

### RNA-Seq data analysis

RNA-seq paired-end reads (read length 150 base pairs) were aligned to the GRCm38 reference genome using hisat2^75^. The average alignment efficiency was 92.7%. Genes were annotated using GENCODE M22^76^ and quantified using HTSeq^77^. Only protein coding genes were used for further analysis. Differential gene expression was determined using DESeq2^78^. Genes were considered differentially expressed if their absolute log2 fold change values were above 1.5 at FDR-adjusted P values (*Padj*) below 0.01, considering a hyperbolic threshold as described previously^15^.

Previously published NB cell line RNA-Seq data were derived from the Cell Line Explorer of Neuroblastoma data (CLEaN)^79^, available at https://ccgg.ugent.be/shiny/clean/.

### Gene sets and gene set enrichment analysis

Preranked gene set enrichment analyses (GSEA) were performed using the R *fgsea* package (*fgseaMultilevel* function, default parameters) with ranking based on the DEseq2 statistic. Mouse and human gene sets were downloaded from the Molecular Signatures Database (MSigDB) v2023.1^80^. ATM, ATR, mTOR and DNAPK related genes were defined based on their presence in the corresponding gene sets as derived from the Pathway Interaction Database (PID), i.e., *PID_ATR_PATHWAY, PID_ATM_PATHWAY, PID_MTOR_4PATHWAY, PID_DNA_PK_PATHWAY*.

### Whole-genome bisulfite sequencing (WGBS)

DNA samples were isolated using DNeasy Kit (QIAGEN) according to standard protocol. The isolated DNA was then subjected to bisulfite conversion and DNA libraries were constructed using a WGBS library preparation kit. Sequencing of the libraries was performed on a next-generation sequencing platform (Illumina) using a paired-end sequencing protocol. The obtained sequencing reads underwent quality control checks, including trimming of adapters, removal of low-quality reads, and elimination of PCR duplicates. Processed reads were then aligned to reference genome (Mouse Grcm38/mm10) and subsequent data analysis was performed to identify differentially methylated regions, visualize methylation patterns, and correlate analyses with other genomic features.

### FACS analysis of immune cells

*Alk-F1178SKI/0;TH-MYCNTg/0* (total n = 14, mixed sex) were, as previously described, followed by ultrasound until the tumour reached a volume of 50-80 mm^3^ (vehicle treated) or 250-400 mm^3^ (elimusertib and lorlatinib treated). They were then treated with either vehicle control or elimusertib (25mg/kg) twice per day for 3 days. Excised tumours were harvested and dissociated with Tumour Dissociation Kit (130-096-730, Miltenyi Biotec B.V. & Co. KG, Bergish Gladbach, Germany) according to the manufacturers instructions. Isolated cells were incubated with antibodies reactive to CD45 (FITC: order # 103108), CD49b (PE; 103506), NKp46 (PECy7; 137618), CD62L (APC; 104412), Gr1 (APCCy7; 108423), CD4 (BV421; 100543), CD8 (BV711; 100747) (Biolegend, San Diego, CA); TCRαβ (PE-CF594; 562841), CD11b (BV510; 562950), CD3 (BV605; 563004), CD11c (BV650; 564079), CD44 (BV786; 563736), F4/80 (BUV395; 565614), B220 (BUV496; 612950), CD69 (BUV737; 612793) (BD Biosciences; Franklin Lakes, NJ); MHCII (AF700; 56-5321-80) (Thermo Fisher Scientific, Waltham, MA) and 7AAD (#11397; Cayman Chemical Company, Ann Arbor, MI) in staining buffer (0.1% BSA in PBS) containing Fc-block (anti-CD16/32; clone 2.4G2), Rat serum and Brilliant stain buffer (563794; BD Biosciences) at 4° C for 60 minutes before washing in PBS two times. Data from cells were analysed on a BD LSRFortessa, and the data subsequently analysed using FlowJo v10 (BD Biosciences).

### CD8-depletion and cGAS/STING-pathway inhibition in mice

*Alk-F1178SKI/0;TH-MYCNTg/0* (n = 20, mixed sex) were screened by ultrasound as previously described, until the tumour reached a volume of 150-300 mm^3^. Treatment was started on day 0 with either anti-mouse CD8α or rat IgG2b isotype control (intraperitoneal injection, 100 µl, 2 mg/kg, diluted in InVivo Pure pH 7.0 Dilution buffer (BioXcell, Catalog #IP0070)). H-151 was delivered by intraperitoneal injection (100 µl, 10mg/kg diluted in 0.1% Tween-80 in PBS). Intraperitoneal injection was repeated on day 3, 7 and 10 for CD8α and IgG2b isotype control while H-151 was given daily between Day 0 and Day 14. ATRi/ALKi (elimusertib 25 mg/kg, lorlatinib 10 mg/kg) combination treatment, as previously described, was initiated on day 1 and continued until day 14. All mice that reached day 14 received all the scheduled intraperitoneal injections except one mouse in the IgG2b isotype control arm who did not receive intraperitoneal injection at day 10. Volume determination was performed on day 0, 7 and 14 and tumour volume quantified as previously described.

### Cell differentiation assays

SK-N-BE(2) cells (6000-8000 cells/well) were seeded in 24-well plates (Eppendorf) overnight and prior to treatment with retinoic acid (RA)(10 μM) or/and elimusertib (10, 20 nM) for 24h. Cell differentiation was recorded in an Incucyte S3 (Essen BioScience). A total of 72 random fields (24 random fields from three independent biological replicates) were analysed by NeuroTrack software (Essen BioScience) which calculates the neurite length (mm/mm^2^) and neurite branch points (mm^2^) automatically. Differentiated cells were counted as a cell with neurite projecting at least 1.5 times of the cell body size. For immunoblotting, SK-N-BE(2) (1 × 10^5^ cells/well) were seeded in 6-well plates overnight, prior to treatment with retinoic acid (RA)(10 μM) or/and elimusertib (10 nM) for 24 h. Primary antibodies against MAP2A (Cell Signalling Technology, #8707), NSE (Abcam, #ab53025), DLG2 (Cell Signalling Technology, #19046), and RET (Cell Signalling Technology, #3220) were employed.

### Statistical analysis

Statistical analyses were performed with either GraphPad Prism 7/8 software or R statistical package (v4.0). Statistical tests are indicated in the respective sections and figure captions. Multiple testing corrections were performed using the Benjamini-Hochberg method.

### Data and code availability

The mass spectrometry proteomics data have been deposited to the ProteomeXchange Consortium^81^ via the PRIDE^82^ partner repository with the dataset identifier PXD041824 and the MassIVE partner repository with the dataset identifier MSV000092789. RNA-seq data are available in ArrayExpress (https://www.ebi.ac.uk/arrayexpress/, accession number: E-MTAB-12961). All other data required to evaluate the conclusions in the paper are provided in Supplementary Materials. Source code used for the RNA-Seq and downstream proteomics analysis is available at GitHub https://github.com/CCGGlab/ATR_AZD.

## Acknowledgements

The authors thank members of the Palmer and Hallberg labs for critical feedback on the manuscript. For proteomics and phosphoproteomics analysis we thank Dr. Egor Vorontsov and Dr. Carina Sihlbom at the Proteomics Core Facility at Sahlgrenska Academy, University of Gothenburg, Sweden. This work has been supported by grants from the Swedish Cancer Society (RHP CAN21/01549; BH CAN21/1525), the Swedish Children’s Cancer Foundation (RHP 2019-0078; BH 2021-0027), the Swedish Research Council (RHP 2019-03914; BH 2021-1192), the Swedish Foundation for Strategic Research (RB13-0204), the Göran Gustafsson Foundation (RHP2016), the Knut and Alice Wallenberg Foundation (KAW 2015.0144), the Assar Gabrielsson’s Foundation (MB FB22-89), the Wenner-Gren Foundation (RHP 2021-0004) and a Ghent University Special Research Fund Starting Grant (JVdE BOF.STG.2019.0073.01).

## Author contributions

B.H., J.V.d.E. and R.H.P. supervised the research project. Wet lab experiments were conducted by G.U., J.G., W.-Y.L. and J.J.. J.V.d.E. and J.L.G. performed RNA-Seq bioinformatics analyses. J.V.d.E., A.S., R.H.P. and J.L.G. performed phosphoproteomics bioinformatics analyses. J.J. performed DNA methylation analyses. D.L., M.B., Y.K. and J.J. performed mouse tumour treatments and monitoring. D.L., M.B. and E.J. performed tumour immunohistochemistry. D.L., A.S. and M.B. performed mouse immune system FACS analyses. R.H.P, A.S., J.G. and J.V.d.E. analysed mass spectrometry proteomics data. R.H.P. coordinated the manuscript writing that was developed with all authors.

## Suppl. figure and table legends

**Figure S1. Transcriptomic response of different NB cell line alterations in DNA damage and related gene sets.**

Bar plot showing RNA-Seq based log2FC values (mean ± 95% confidence interval) of 276 genes involved in the DNA Damage Response, Hallmark G2M checkpoints of Hallmark E2F targets as indicated on top and for different neuroblastoma cell lines and interventions as indicated on the right and left respectively. Cell line data derived from CLEaN^79^.

**Figure S2. Selected phosphorylation response of NB cell lines to ATR inhibition.**

CLB-BAR NB cells were synchronized by thymidine block and treated with either DMSO (Ctrl) or 1 µM ceralasertib for 6 or 12 h, as indicated. Lysates were immunoblotted with anti-pATR, pATM, pFOXM1 and pCHK1. Tubulin was used as a loading control. NSC, non-synchronized cells; SC, synchronized cells.

**Figure S3. Phosphoproteomic response of NB cell lines to ATR inhibition.**

CLB-BAR cells were synchronized by thymidine block and treated with either DMSO (control), elimusertib (50 nM) and ceralasertib (50 nM or 1 µM) for 6 h. **A.** Volcano plots showing differential phosphorylation (DP) between different treatments (as indicated on top) and control conditions. ATM, ATR, DNAPK and RPS6 sites (indicated on the left) shown in blue. **B.** Bar plots showing the mean +/- 95% confidence intervals from all sites for the different treatments and clusters (see heatmap shown in main Fig. 4E) as indicated.

**Figure S4. Flow cytometric analysis of immune cells in tumour tissues.** Mice with tumours were treated with elimuseratib or lorlatinib for three days or left without treatment (control) and single cells were isolated from tumour tissues. **A.** Equal numbers of CD45^+^ cells from individual mice were concatenated and were visualized in two dimensions using tSNE. Individual cells are coloured according to treatment group are shown to the left. Based on expression of TCRαβ, CD4, CD8α, GR1, NKp46, CD49, B220 and MHCII major immune cell types were identified (right). **B.** Bar graphs were constructed that show the proportion of CD45+ cells belonging to the different types in individual mice with the bar indicating the mean. **C-E.** T cells were further analysed based on them being CD4 or CD8 expressing **(C),** whether they belonged to CD62L^+^CD44^-^ naïve, CD62L^+^CD44^+^ central memory (CM), CD62L^-^ CD44^+^ effector memory (EM) or CD62L^-^CD44^-^ subsets **(D),** or to visualise expression of CD69 in individual mice **(E)**. **F-H.** Cells belonging to the MonoMacroDC subset were visualised using tSNE to identify nine subsets of cells. Left hand graph indicates subsets identified based on expression of CD45, MHCII, CD11b, CD11c, GR1, F4/80, CD8α and B220 (graph) **(F)**. Phenotypic comparison of the nine subsets **(G)** and the proportion of MonoMacroDC belonging to each cell type **(H)** in individual mice. The data was collected on two experimental dates with a total of six elimuseratib treated, three lorlatinib treated and 5 control mice. Statistical significance was calculated using one-way ANOVA with Tukey’s multiple comparison test (* p <0.05, **p < 0.01).

**Figure S5. Chemical and Genetic Perturbations gene set enrichment analysis (GSEA) on elimusertib-treated tumours.** Volcano plots showing GSEA results using known chemical and genetic perturbations gene sets from MSigDB. Enrichment was performed using results from DGE between NB tumours from *Alk-F1178S;Th-MYCN* mice in untreated control conditions (n=6) and after 3 days of elimusertib treatment (n=3). Neuroendocrine and neuron-related gene sets indicated. Bottom plots showing running score plots for as indicated.

**Figure S6. Elimusertib treated tumours exhibit a robust methylation response A.** Violin plot showing levels of differentially methylated regions (DMRs) between vehicle and elimusertib treated *Alk-F1178S;Th-MYCN* tumours. **B.** Diagram depicting DMR length distribution in elimusertib treated *Alk-F1178S;Th-MYCN* tumours. **C.** Bar graph showing numbers of hypo- and hypermethylated DMRs within functional genomic regions. **D.** Diagram showing DMR levels within, upstream, and downstream of gene bodies in elimusertib treated *Alk-F1178S;Th-MYCN* tumours compared to vehicle treated controls. All DMRs were sorted by statistic areaStat including hypomethylated (areaStat<0) and hypermethylated (areaStat>0) DMRs.

**Table S1. Phosphoproteomics differential phosphorylation results.** CLB-BAR NB cell lines were treated for 6 hours with elimusertib (50 nM) or ceralasertib (50 nM & 1µM) as indicated by tab names.

**Table S2. RNA-Seq differential expression results.** Mice were treated for 3 days with 25 mg/kg elimusertib or ceralasertib as indicated by tab names.

